# PAM-flexible adenine base editing rescues hearing loss in a humanized *MPZL2* mouse model harboring an East Asian founder mutation

**DOI:** 10.1101/2024.10.29.620803

**Authors:** Shao Wei Hu, Sohyang Jeong, Luoying Jiang, Hansol Koo, Zijing Wang, Won Hoon Choi, Biyun Zhu, Heeyoung Seok, Yi Zhou, Min Gu Kim, Dan Mu, Huixia Guo, Ziyi Zhou, Sung Ho Jung, Yingting Zhang, Ho Byung Chae, Liheng Chen, Sung-Yeon Lee, Luo Guo, Myung-Whan Suh, Yang Xiao, Moo Kyun Park, Honghai Tang, Jae-Jin Song, Xi Chen, Ai Chen, Jun Ho Lee, Sangsu Bae, Sang-Yeon Lee, Yilai Shu

**Author notes:** To whom correspondence should be addressed to S.B., S-Y.L., and Y.S. These authors contributed equally (Shao Wei Hu, Sohyang Jeong, Luoying Jiang, Hansol Koo).

## Abstract

Hearing loss is one of the most prevalent sensory disorders, but no commercial biological treatments are currently available. Here, we identified an East Asia-specific founder mutation, the homozygous c.220C>T mutation in *MPZL2*, that contributes to a significant proportion of hereditary deafness cases in our cohort study. We found that the disease-causing mutation could be targetable by adenine base editors (ABEs) that enable A·T-to-G·C base corrections without DNA double-strand breaks. To demonstrate this, we developed a humanized mouse model (*hMPZL2*^Q74X/Q74X^) that recapitulates human *MPZL2* deafness and leads to progressive hearing loss. A PAM-flexible ABE variant with reduced bystander and off-target effects (ABE8eWQ-SpRY:sgRNA3) was packaged in dual adeno-associated viruses (AAVs) and injected into the inner ear of *hMPZL2*^Q74X/Q74X^ mice and effectively corrected the mutation. This treatment significantly restored hearing function, improved inner ear structural integrity, and reversed altered gene expression. Base editing may hold therapeutic potential for hereditary deafness, including most cases of *MPZL2* deafness.

## Introduction

Hearing loss is the most common sensory disorder in humans and affects about 466 million people worldwide (World Health Organization, https://who.int/news-room/fact-sheets/detail/deafness-and-hearing-loss). Although no current treatments are capable of fully restoring biological hearing function^1^, advancements in our understanding of the genetic architecture and molecular mechanisms of hearing loss have led to significant progress in inner ear gene therapies^2,3^. Notably, CRISPR genome editing technologies have been harnessed to directly correct disease-causing mutations for the fundamental treatment of genetic disorders^4^. In particular, base editors, including cytosine base editors (CBEs) and adenine base editors (ABEs), are engineered genome-editing tools that enable the precise repair of targeted genomic base pairs without double-strand breaks^5,6^. Presently, ABEs and CBEs have been successfully used in treating hearing loss in hereditary deafness mouse models^7,8^, suggesting that base editors hold potential as one-time treatments for hereditary deafness caused by pathogenic point mutations.

Recessive loss-of-function mutations represent approximately 80% of hereditary deafness cases^9^. One of these, non-syndromic autosomal recessive deafness-111 (DFNB111), is a leading cause of mild-to-moderate progressive hearing loss^10^. Here, we identified an East Asia-specific founder mutation of DFNB111, the homozygous c.220C>T variant in *MPZL2*, that contributes to a significant proportion of hereditary deafness cases in our cohort study. To model DFNB111, we generated a humanized *MPZL2* c.220C>T knock-in mouse model that replicates the human condition. For *in vivo* editing, we tested various combinations of ABE variants, four types of Cas9 variants with different PAMs, and single-guide RNA (sgRNA)s to correct the c.220C>T mutation. Ultimately, we determined the optimal PAM-flexible ABE variant (i.e., ABE8eWQ-SpRY:sgRNA3), which achieved a 60% editing efficiency *in vitro* with no detectable bystander or off-target effects. The administration into the inner ear of ABE8eWQ-SpRY:sgRNA3 packaged in dual AAV-ie vectors successfully restored both auditory function and the structural integrity of the inner ear. The development of humanized mouse models and the successful correction of the *MPZL2* founder mutation using a single PAM-flexible ABE together represent a significant advancement in base editor gene therapy, suggesting great promise for treating hereditary deafness, including most DFNB111 cases.

## Results

### Clinical significance of the East Asia-specific founder mutation in *MPZL2*-related DFNB111

We reviewed our sensorineural hearing loss (SNHL) cohort, which consisted of 1,437 unrelated families attending the hereditary deafness clinic within the Otorhinolaryngology Division at two tertiary centers (Seoul National University Hospital (SNUH) and the Eye & ENT Hospital, Fudan University (EENT)), between March 2021 and February 2024. Within this cohort, a filtering process yielded 234 pediatric probands (≤18 years old) with symmetric, mild-to-moderate (range 21–55 dB hearing threshold), non-syndromic SNHL **(Fig.1a)**. Through comprehensive genetic testing using exome/genome sequencing, we identified disease-causing mutations in 155 probands (65.7%). The detailed genotypes and their pathogenicity are described in **Supplementary Table 1**. Collectively, 17 deafness genes that were seen in 2 or more probands were identified as disease-causing within these 155 genetically diagnosed families. *GJB2* was the most frequently affected gene (36.8%, 57/155), followed by *STRC* (18.1%, 28/155) and *MPZL2* (9.0%, 14/155) (**Fig.1b**). In our cohort, we identified a total of 24 *MPZL2*-associated DFNB111 patients from 20 unrelated families, regardless of age at ascertainment. The pedigrees, genotypes, and hearing loss phenotypes are presented in **Figure 1c–d**. The significant contribution of the *MPZL2* c.220C>T mutation to DFNB111 was found. This C→T mutation at position c.220 (c.220 C ⋅G to T⋅A) creates a stop codon (TAG) that replaces the Gln codon (CAG) at p.74 (Q74X), likely resulting in nonsense-meditated mRNA decay or a truncated protein. In our DFNB111 patients, 23 (95.8%) harbored at least one c.220C>T allele. Notably, 19 of these patients (79.2%) were homozygous for the c.220C>T mutation (**Fig.1e**). The mutational landscape of DFNB111, as documented in the literature and including our cohort, is illustrated in **Extended Data Fig. 1 a**, highlighting the high mutational burden of the c.220C>T mutation. The c.220C>T allele recurred frequently, especially in East Asian populations, suggesting a founder mutation in East Asia **(Fig.1f)**. In contrast, the second most frequently recurrent allele, c.72delC, was found across different genetic ancestries. Auditory function-gene profiles exhibited significant progressive hearing loss over the course of decades (**Fig.1g**), specifically an annual progression of hearing loss of 0.61 dB at low frequencies, 0.57 dB at middle frequencies, and 0.61 dB at high frequencies (**Extended Data Fig. 1b**). Collectively, we hypothesized that a single ABE that corrects the c.220C>T founder mutation could serve as a “one-and-done” therapy, potentially treating or even curing the majority of DFNB111 patients, which constitute a significant proportion of all hereditary deafness cases.

**Figure 1.**
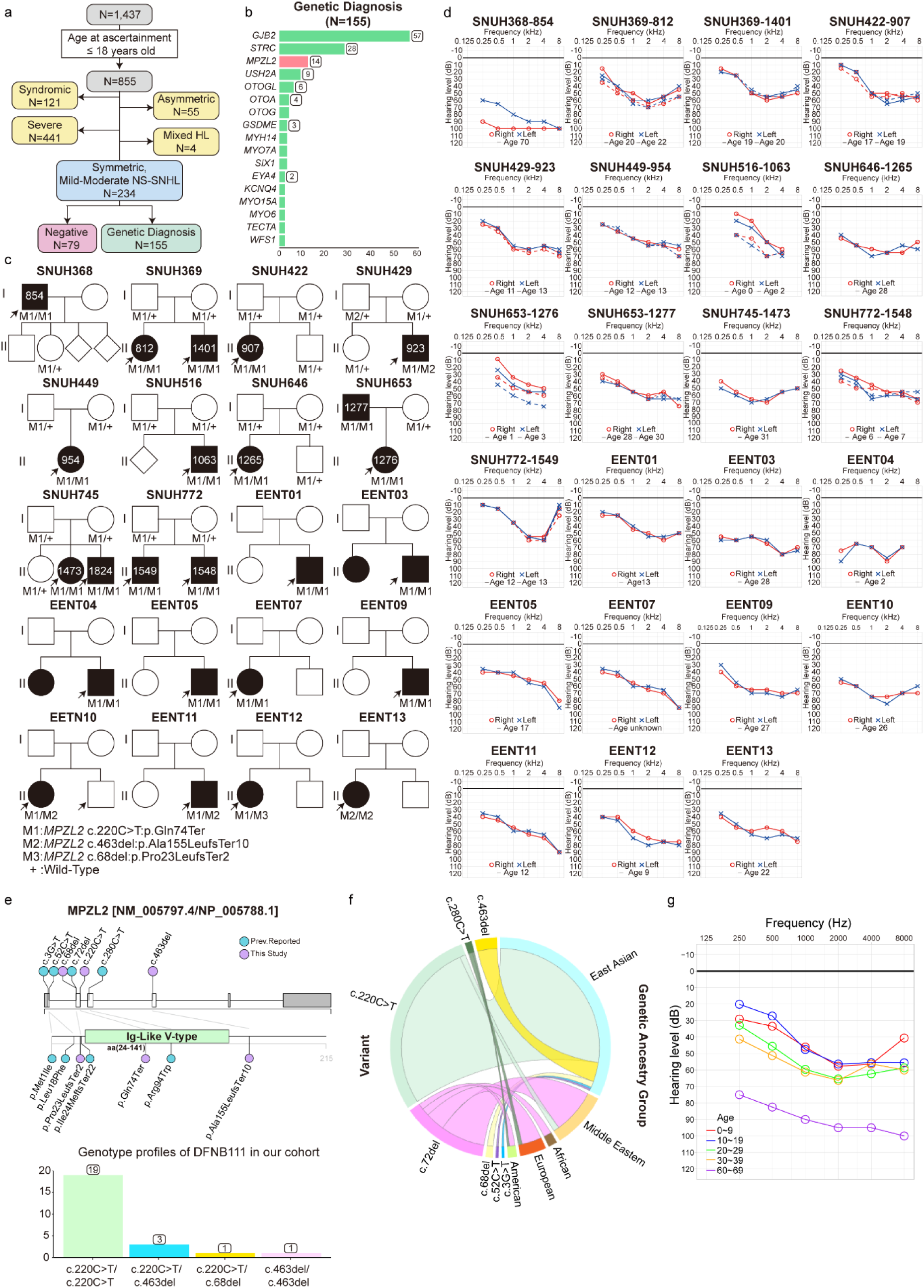
Clinical significance of the *MPZL2* c.220C>T mutation. (a) Schematic flow diagram illustrating the genetic diagnosis study of symmetric, mild-to-moderate, non-syndromic sensorineural hearing loss (ns-SNHL) in children from two tertiary centers. (b) Gene signatures of symmetric, mild-to-moderate, ns-SNHL in children. The bar plot shows the frequencies of 17 deafness genes that were seen in 2 or more probands from 155 genetically diagnosed families. (c) Pedigrees and genotypes of 24 affected patients from 20 unrelated DFNB111 families. Arrows, probands; Black filled circles or rectangles, affected patients. (d) Serial audiograms of 23 affected DFNB111 patients. Red, right ear; Blue, left ear. (e) *MPZL2* mutational landscape on a Lollipop plot (upper), and the prevalence of *MPZL2 in trans* mutation combinations in our cohort (lower). (f) The mutational burden of different *MPZL2* mutations depending on the genetic ancestry groups in the Chord diagram. (g) The natural course of hearing loss in DFNB111 patients over decades across different hearing frequencies.

### Generation and characterization of the *MPZL2* humanized mouse models

Because the mouse *Mpzl2* and human *MPZL2* sequences differ near the c.220C>T mutation, we decided to develop a humanized knock-in mouse model of DFNB111 by inserting 648 bp of human *MPZL2* complementary DNA (cDNA) containing the pathogenic c.220C>T mutation into the mouse *Mpzl2* locus (**Fig. 2a**). The integration of the human cassette and successful on-targeting was verified (**Extended Data Fig.2a**). Additionally, the sequence of the *hMPZL2*^Q74X^ mice was confirmed using Sanger sequencing (**Fig.2b**). In the humanized knock-in mouse model of DFNB111 (*hMPZL2*^Q74X/Q74X^), the mRNA transcript levels of human *MPZL2* and mouse *Mpzl2* were undetectable across tissues, in contrast to *Mpzl2* wild-type (WT) mice (**Fig.2c**). Consistent with previous studies^11,12^, we found that Mpzl2 protein was expressed in the organ of Corti, primarily in outer hair cells (OHCs), Deiters’ cells (DCs), and at the contact between DCs and the basilar membrane during both the neonatal period (P4) and the adult period (P28) in *Mpzl2*^WT^ mice **(Extended Data Fig.2b, c)**. Conversely, the MPZL2/Mpzl2 protein expression was absent in *hMPZL2*^Q74X/Q74X^ mice at both P4 and P28. Thus, the c.220C>T mutation (Q74X) is interpreted to cause nonsense-meditated mRNA decay rather than producing a truncated protein. At 4 weeks of age, *hMPZL2*^Q74X/Q74X^ mice exhibited an elevation of 14.5 dB at 24 kHz and 23.5 dB at 32 kHz in auditory brainstem response (ABR) thresholds compared to *Mpzl2*^WT^ mice (**Fig.2d**), indicating the onset of hearing loss beginning at 4 weeks, specifically at higher frequencies. Similar to an *Mpzl2* knock-out mouse model^12^, *hMPZL2*^Q74X/Q74X^ mice initially displayed mild-to-moderate hearing loss, which progressively worsened to severe hearing loss. At 12 weeks of age, the ABR thresholds in *hMPZL2*^Q74X/Q74X^ mice exceeded 70 dB across the click and 4–32 kHz frequencies, except at 8 kHz. In contrast, *hMPZL2*^Q74X/WT^ mice retained ABR thresholds comparable to *Mpzl2*^WT^ mice (**Fig. 2d**). We also performed distortion product otoacoustic emission (DPOAE) measurements to assess the function of OHCs. Similarly, DPOAE thresholds in *hMPZL2*^Q74X/Q74X^ mice were significantly higher than those in *hMPZL2*^Q74X/WT^ and *Mpzl2*^WT^ mice (**Fig.2e**), suggesting that the *MPZL2* c.220C>T mutation reduces the functionality of OHCs. In *hMPZL2*^Q74X/Q74X^ mice, a significant loss of OHCs and DCs in the organ of Corti was observed, particularly in the mid- and high-frequency regions of the cochlea, along with cellular disarrangement and a collapsed tunnel of Corti (**Fig. 2f and Extended Data Fig.3**). In contrast, no distinct structural abnormalities were observed in the stria vascularis or spiral ligament, and the density of spiral ganglion neurons (SGNs) remained unaffected (**Extended Data Fig.3,4**). To evaluate whether the hearing loss in *hMPZL2*^Q74X/Q74X^ mice is affected by the insertion of human cDNA into the mouse *Mpzl2* locus, we also generated another humanized *MPZL2*^WT^ mouse model (*hMPZL2*^WT/WT^) using a sequential knock-in strategy (**Extended Data Fig.5a**), and we verified the successful on-targeting and sequence of *MPZL2*^WT^ (**Extended Data Fig.5b, c**). The *hMPZL2*^WT/WT^ mice exhibited normal thresholds across frequencies at 12 weeks of age, comparable to *Mpzl2*^WT^ mice (**Extended Data Fig.6a, b**), and displayed normal inner ear structures (**Extended Data Fig.6c, d**). These observations indicate that the auditory and inner ear phenotypes in hMPZL2^Q74X/Q74X^ mice are specifically caused by the pathogenic c.220C>T mutation.

**Figure 2.**
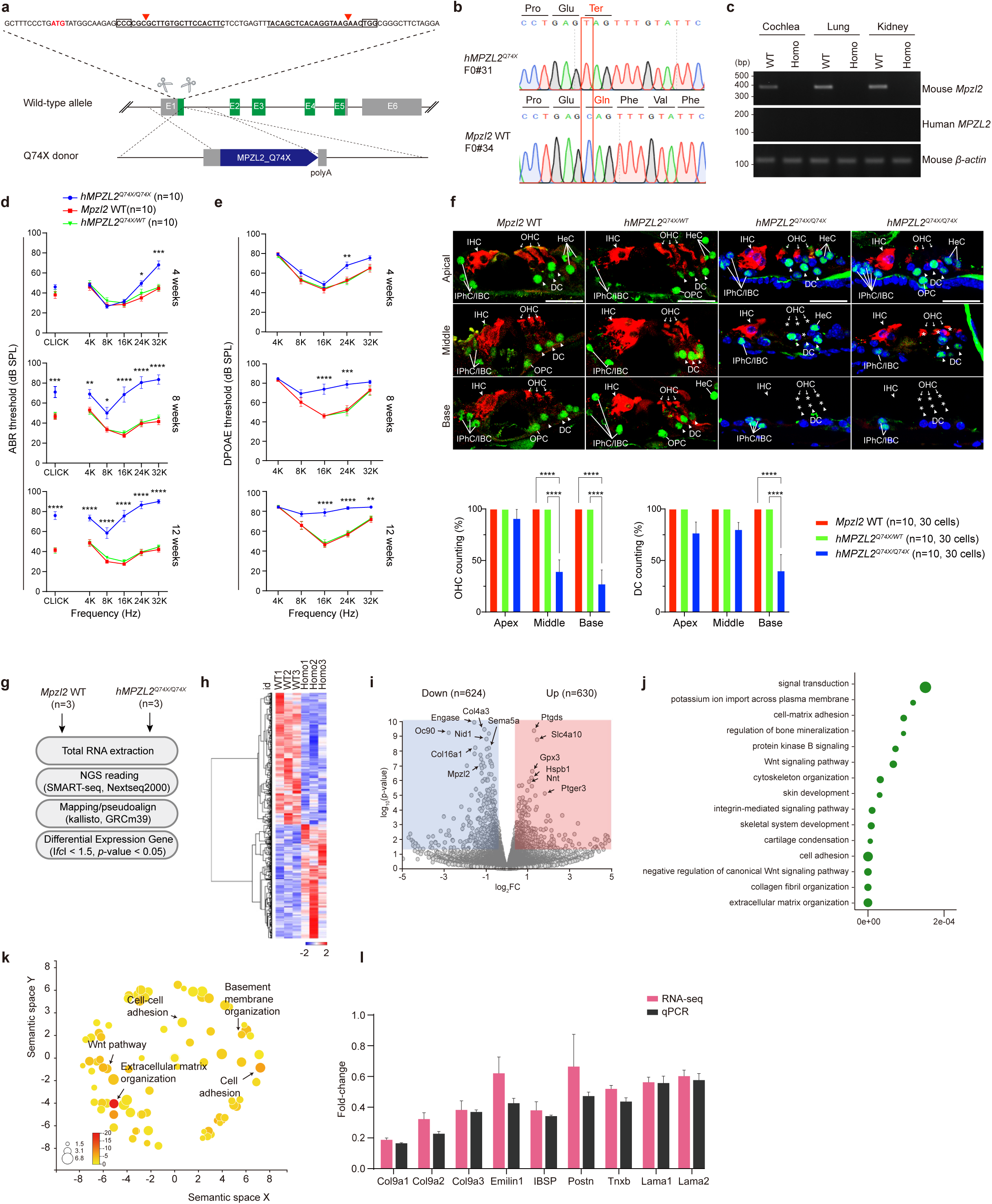
Generation and characterization of humanized mouse models. (a) Targeting strategy for the humanized *MPZL2* mouse model using the CRISPR/Cas9 system. The *hMPZL2*^Q74X^ mouse was generated by microinjection of a human *MPZL2*^Q74X^ expression cassette using a dsDNA donor. The black boxes and lines indicate the positions of the sgRNAs with PAMs, and the red letters and red arrowheads indicate the start and stop codons and the Cas9 cleavage sites, respectively. (b) Sequence verification of *hMPZL2*^Q74X^ mice and *Mpzl2*^WT^ mice using Sanger sequencing. (c) RT-PCR analysis to detect *MPZL2* and *Mpzl2* expression in P4 and *P28 hMPZL2*^Q74X/Q74X^ and *Mpzl2*^WT^ mice. (d) Comparison of ABR thresholds among *Mpzl2*^WT^ (red, n = 10), *hMPZL2*^Q74X/WT^ (green, n = 10), and *hMPZL2*^Q74X/Q74X^ (blue, n = 10) mice at 4, 8, and 12 weeks of age. ABR thresholds were recorded in response to click and 4–32 kHz tone bursts. (e) DPOAE thresholds in *hMPZL2*^Q74X/Q74X^ mice (blue, n = 10) compared with *Mpzl2*^WT^ mice (red, n = 10) and *hMPZL2*^Q74X/WT^ mice (green, n = 10) at 4, 8, and 12 weeks of age. Data are presented as the mean ± SEM. Statistical significance was determined using one-way ANOVA with Bonferroni’s multiple comparisons tests. Significance levels are indicated as \**P* < 0.05, \*\**P* < 0.01, *** *P* < 0.001, and \*\*\*\**P* < 0.0001. (f) Representative section images (10 μm) from 12-week-old *Mpzl2*^WT^ mice (n = 1), *hMPZL2*^WT/Q74X^ mice (n = 1), and *hMPZL2*^Q74X/Q74X^ mice (n = 2). Sections were immunolabeled with Myosin VIIa (red) to mark inner hair cells (IHCs) and outer hair cells (OHCs) and with Sox2 (green) to identify supporting cells (SCs), including Deiter’s cells (DCs), in the apical, middle, and basal turns of the organ of Corti. Arrowheads indicate IHCs and DCs, white arrows point to OHCs, and the white line marks SCs. Asterisks indicate loss of OHCs and DCs. Scale bar: 25 μm. OHCs and DCs were counted in *Mpzl2*^WT^ mice (n = 10; red), *hMPZL2*^Q74X/WT^ mice (n = 10; green), and *hMPZL2*^Q74X/Q74X^ mice (n = 10; blue). Data are shown as the mean ± SEM. Statistical significance was determined using one-way ANOVA with Bonferroni’s multiple comparisons tests, and significance levels are indicated as \**P* < 0.05, \*\**P* < 0.01, \*\*\**P* < 0.001, and \*\*\*\**P* < 0.0001. (g) Schematic diagram of the RNA-sequencing analysis. Each group consisted of three biological replicates (n = 3). (h) Heatmap analysis of DEGs (upregulation, red; downregulation, blue). (i) Volcano plot of DEGs. The numbers of upregulated (red box) and downregulated (blue box) genes are indicated. (j) Gene ontology (GO) enrichment analysis in biological processes (BP). (k) Revigo visualization of the top 20 GO BP terms associated with cell adhesion, basement membrane organization, Wnt pathway, and ECM organization. The size of the circle indicates the relative number of genes in each gene set, and the color indicates the significance. (l) Quantitative RT-PCR assay for validating the DEGs of interest, *COL9A1/2/3, EMILIN1, IBSP, POSTN, TNXB, LAMA1/2*. Data are presented as the mean ± SEM.

We subsequently conducted RNA-seq analysis to explore the molecular pathways involved in the cochlear histopathology caused by the c.220C>T mutation. Cochlear membranous labyrinth tissue from *hMPZL2*^Q74X/Q74X^ and *Mpzl2*^WT^ mice was dissected at P28 and used for RNA-seq analysis **(Fig.2g)**. Principal Component Analysis of the total RNA-seq data revealed a distinct clustered pattern of gene expression, and a heatmap visualized the patterns and clusters of individual variable values between *hMPZL2*^Q74X/Q74X^ and *Mpzl2*^WT^ mice **(Fig.2h).** We identified a total of 1,254 differentially expressed genes (DEGs) **(Fig.2i)**, and Gene Ontology (GO) analysis of the DEGs revealed significant enrichment in biological process, cellular component, and molecular function terms **(Fig. 2j and Extended Data Fig.7)**. REVIGO analysis further highlighted significant enrichment in biological processes related to cell adhesion, extracellular matrix (ECM) organization, and basement membrane integrity **(Fig.2k).** We filtered the DEGs using the gEAR (http://umgear.org) and SHIELD databases (http://shield.hms.harvard.edu), and we further refined the list to focus on DEGs associated with cell adhesion, ECM organization, and basement membrane integrity **(Supplementary Table 2).** The associated transcripts, including *COL9A1/2/3*, *EMILIN1, IBSP, POSTN, TNXB,* and *LAMA1/2*, identified through the RNA-seq analysis were validated by quantitative RT-PCR **(Fig.2l)**. Collectively, the RNA-seq results suggested that the *MPZL2* c.220C>T mutation disrupts cellular pathways involved in cell adhesion, ECM organization, and basement membrane integrity, thus contributing to the histopathological changes observed in *hMPZL2*^Q74X/Q74X^ mice.

### In vitro selection of the optimal ABE and sgRNA combination to correct the *MPZL2* c.220C>T mutation

To select the best base editing system *in vitro*, we constructed a HEK293T cell line that mimics the pathology of human DFNB111 patients by introducing a nucleotide change (c.220C>T) at the *MPZL2* genome locus, resulting in a nonsense mutation (p.Q74X), and used it as a model for base editor screening (**Fig.3a**). For successful ABE-based base editing, an appropriate PAM is required to position the target base within the optimal editing window, approximately 3–8 positions from the 5’ end of the protospacer. Thus, we searched for an NGG PAM sequence that the nCas9 of ABE could recognize near the target adenine. However, the target site only had one sgRNA binding site with the only available NGG PAM, which placed the target adenine at spacer position 2 (i.e., A_2_), which was outside the editing window where ABE works best (**Fig.3b**). Given the lack of suitable NGG PAMs, we explored the use of engineered or evolved Cas9 nuclease-based ABE variants targeting non-NGG PAM. Thus, we included four types of Cas9 variants with different PAM requirements: NG-Cas9^13^ (NG), SpRY^14^ (NNN; NAN, NTN, and NCN), and eNme2-C^15^(N4NC) in addition to the canonical SpCas9^16^ (NGG). Moreover, we also considered different ABE platforms including ABEmax, ABE8eWQ, and ABE8e^17^, which exhibited different editing activities and different editing windows. Consequently, we reconstituted a total of 14 ABE:sgRNA combinations using different adenine deaminases and Cas9 variants, and we designed all possible 6 sgRNAs to more optimally target the c.220C>T mutation in the *MPZL2* allele. The target adenine was at position 2, 4, 5, 6 and 7 with the corresponding AGG, AG, TAG, CTA, TCT, and AGGACG PAMs, and these were referred to as sgRNA1–6. All ABE:sgRNA combinations were co-transfected into the HEK293T-*MPZL2* mutant cells, and the on-target editing efficiency was analyzed by targeted deep sequencing. The results showed that SpRY-based ABE with sgRNA3 (NAN; TAG PAM) and sgRNA4 (NTN; CTA PAM) induced significantly higher levels of A-to-G editing at the target adenine compared to the other ABE:sgRNA combinations (**Fig.3c**). We also thoroughly evaluated the A-to-G editing of all bystander adenines. Notably, the combination of ABE8eWQ-SpRY:sgRNA3 showed high precision by inducing less bystander editing and preserving high targeted editing efficiency (Bystander A1: 0.1%, A2: 0.2% and Target A5: 54%, Silent A9: 1%) (**Fig.3d**), whereas ABE8e-SpRY induced base editing at both the target adenine and bystander adenines (Bystander A1: 0.3%, A2: 1.8% and Target A5: 42%, Silent A9: 37%). This indicates that ABE8e had a wider editing window (3–10 bp) than ABE8eWQ (4–8 bp), which is consistent with a previous report^18^. Taken together, our data showed that it would be best to use ABE8eWQ-SpRY and ABEmax-SpRY with sgRNA3 and sgRNA4 to correct the target c.220C>T in exon 2 of *MPZL2*, and we finalized the combination of ABE8eWQ-SpRY:sgRNA3 for further *in vivo* study considering that ABE8eWQ is known to have fewer sgRNA-independent RNA off-target effects compared to ABEmax^19,20^.

### Off-target analysis in the human genome

To evaluate potential sgRNA-dependent off-target reactions in HEK293T-*MPZL2* mutant cells treated with ABE8eWQ-SpRY:sgRNA3, we screened for genome-wide off-target sites for SpRY using Cas-OFFinder software^21^, and we performed a PAMless NNN search that allowed for up to 2 mismatches and/or 1 DNA/RNA bulge outside the seed region at position 10–18 from the 5’ end. We identified 18 potential off-target sites (OT1–OT18) in the human genome (**Fig.3e, Supplementary Table 3**). In all, base editing rates at the off-target sites were similar to controls, but we only detected a slight increase in the A-to-G conversion rate at position 5 at OT10, which is in the exon regions of the *CDC34* gene **(Fig.3f**). However, there have been no clinical phenotypes associated with the *CDC34* gene (https://www.omim.org/). We next evaluated sgRNA-independent RNA off-targets of ABE8eWQ-SpRY compared with two other ABE variants, ABEmax-SpRY and ABE8e-SpRY. We co-transfected each ABE variant along with sgRNA3 into HEK293T-*MPZL2* mutant cells and analyzed the frequency of A-to-I conversion in three representative RNA transcripts (*CCNB1IP1*, *AARS1*, and *TOPORS*)^20,22^. Across all three mRNA transcripts, ABE8eWQ-SpRY induced lower RNA off-target effects compared to ABEmax-SpRY and ABE8e-SpRY, and these were similar to controls **(Fig. 3g, h**). These data further support that ABE8eWQ-SpRY was the best option in our case.

**Figure 3.**
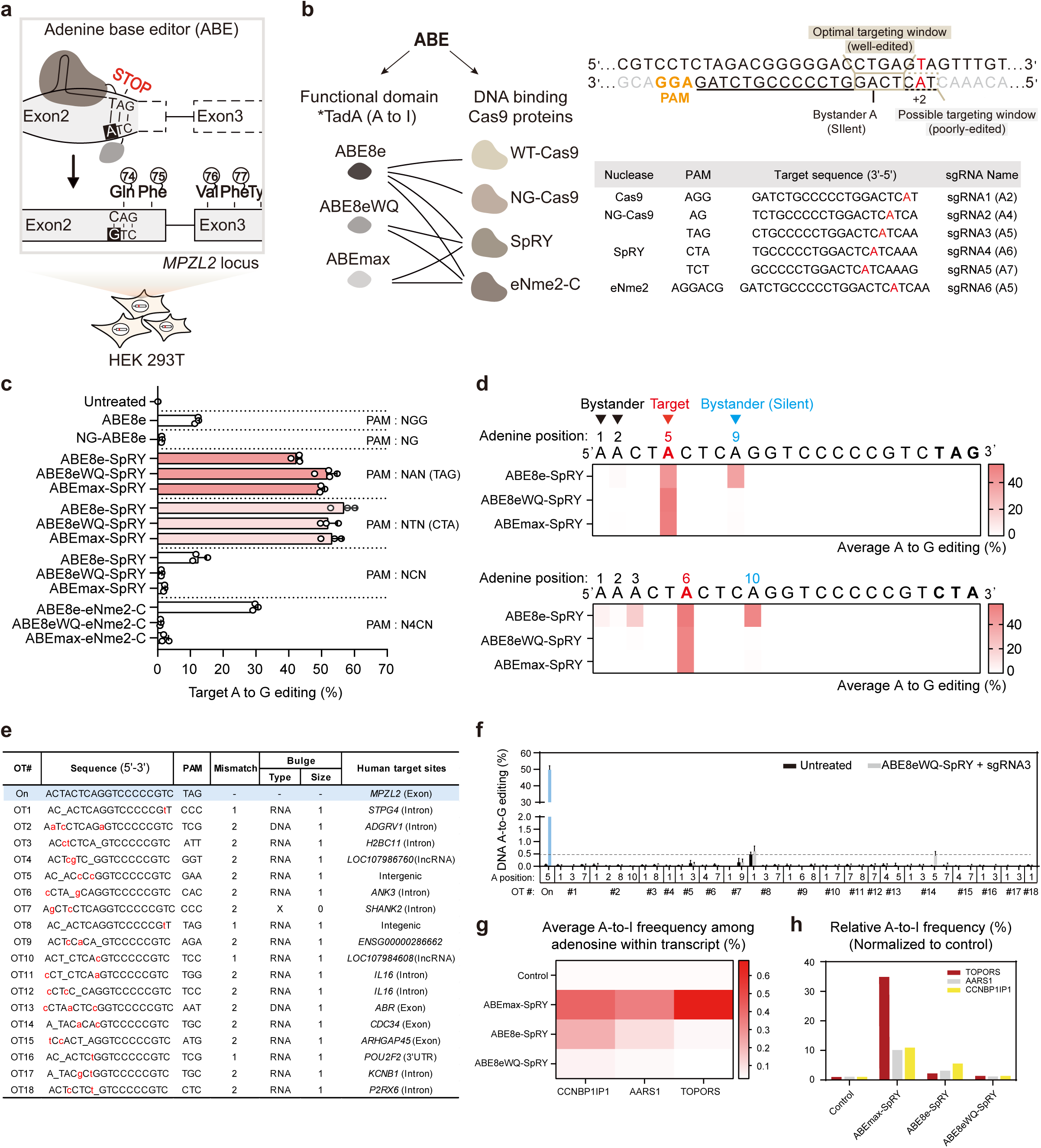
*In vitro* selection of the optimal ABE system for correcting the *MPZL2* c.220C>T mutation. (a) Schematic representation of a correction strategy using PAM-flexible ABEs in HEK293T monoclonal cells harboring the homozygous C>T nonsense mutation (c.220C>T: p.Q74X) at the *MPZL2* locus. (b) Design of sgRNAs targeting the *MPZL2* c.220C>T mutation. For more effective editing, the target adenine A needs to be within ABE’s preferred targeting window. In the table below, 6 sgRNA candidates featuring different PAMs are listed, each placing the target A (red) at different positions on the protospacer: A2, A4, A5, A6, and A7. (c) The comparison of the on-target DNA A-to-G editing efficiencies of all ABEs used in this study. Bars reflect the mean value of A-to-G editing efficiencies on adenine derived from three or four independent experiments. (d) For the selected ABEs based on nSpRY nuclease, the A-to-G conversion rate for each adenine position within the 20-bp protospacer was averaged and shown in a heat map. The numbers at the top of the heatmap indicate adenine positions counted from the 5’ end. Bystander bases are shown as black arrows, silent mutations as blue arrows, and target bases as red arrows. (e) List of potential off-target sites relative to the target site. The potential off-target sites were found using Cas–OFFinder, and we included those with up to two mismatches (excluding an extra G at the 5’ end of the sgRNA) and/or up to one DNA/RNA bulge. (f) A- to-G conversion rates at in silico-predicted off-target sites after ABE8eWQ-SpRY and sgRNA3 treatment. The off-target analysis was conducted on the cells for which on-target analysis had been performed. (g) Targeted RNA off-target effects in HEK-293T cells mimicking the *MPZL2* c.220C>T variant with three SpRY ABEs that were expressed using NAN PAM sgRNA. The average frequency of A-to-I transitions in three mRNA transcripts (*CCNB1IP1*, *AARS1*, and *TOPORS*) with each of the SpRY-mediated ABE variants. (h) The bars indicate the average of the normalized editing efficiencies. The average editing efficiencies of three repeats for each target were normalized to the editing efficiency of the untreated control.

**Figure 4.**
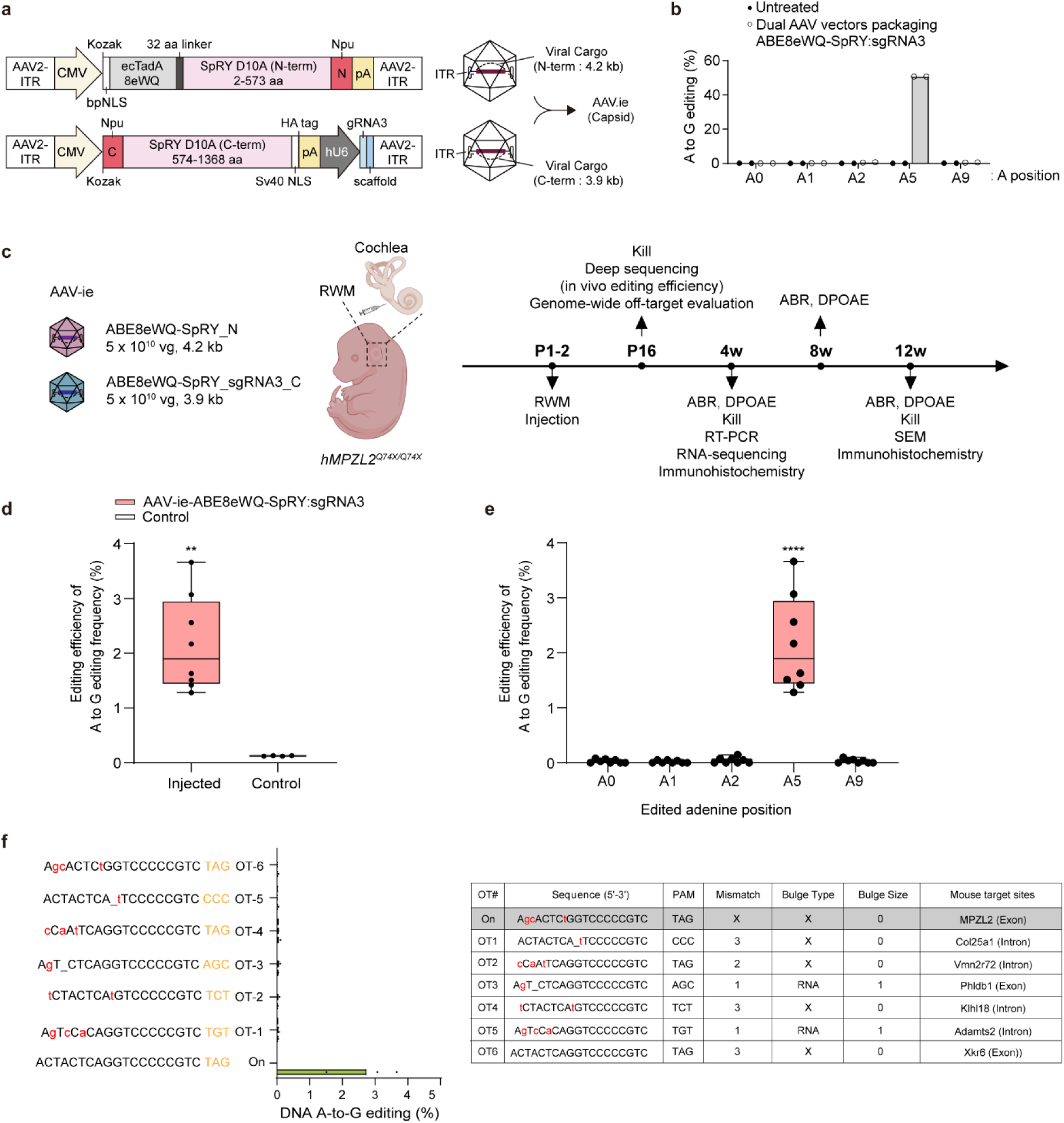
*In vivo* correction of the *MPZL2* c.220C>T mutation by dual AAV-ie-ABE8eWQ-SpRY:sgRNA3. (a) Schematic representation of the dual AAV constructs utilizing split-intein for ABE delivery *in vivo*, resulting in intein-mediated assembly of complete ABE:sgRNA complexes. The dual vectors were packaged into the AAV serotype AAV-ie. (b) Assessment of the editing efficiencies of the *MPZL2* target adenine and other bystander adenines using dual AAV vectors that encoded split-intein ABE8eWQ-SpRY and sgRNA3. (c) Experimental overview of the *in vivo* base editing. (d, e) *In vivo* efficiency of A·T to G·C editing at on-target sites and bystander effects in DNA extracted from the organ of Corti (n = 8). Statistical analyses were performed using Student’s t-test (d) and one-way ANOVA (e). Significance levels are indicated as \**P*< 0.05, \*\**P* < 0.01, \*\*\**P* < 0.001, and \*\*\*\**P* < 0.0001. (f) *In vivo* potential off-target sites relative to the target site. The potential off-target sites were identified using Cas–OFFinder, including those with up to two mismatches and/or up to one DNA/RNA bulge.

**Figure 5.**
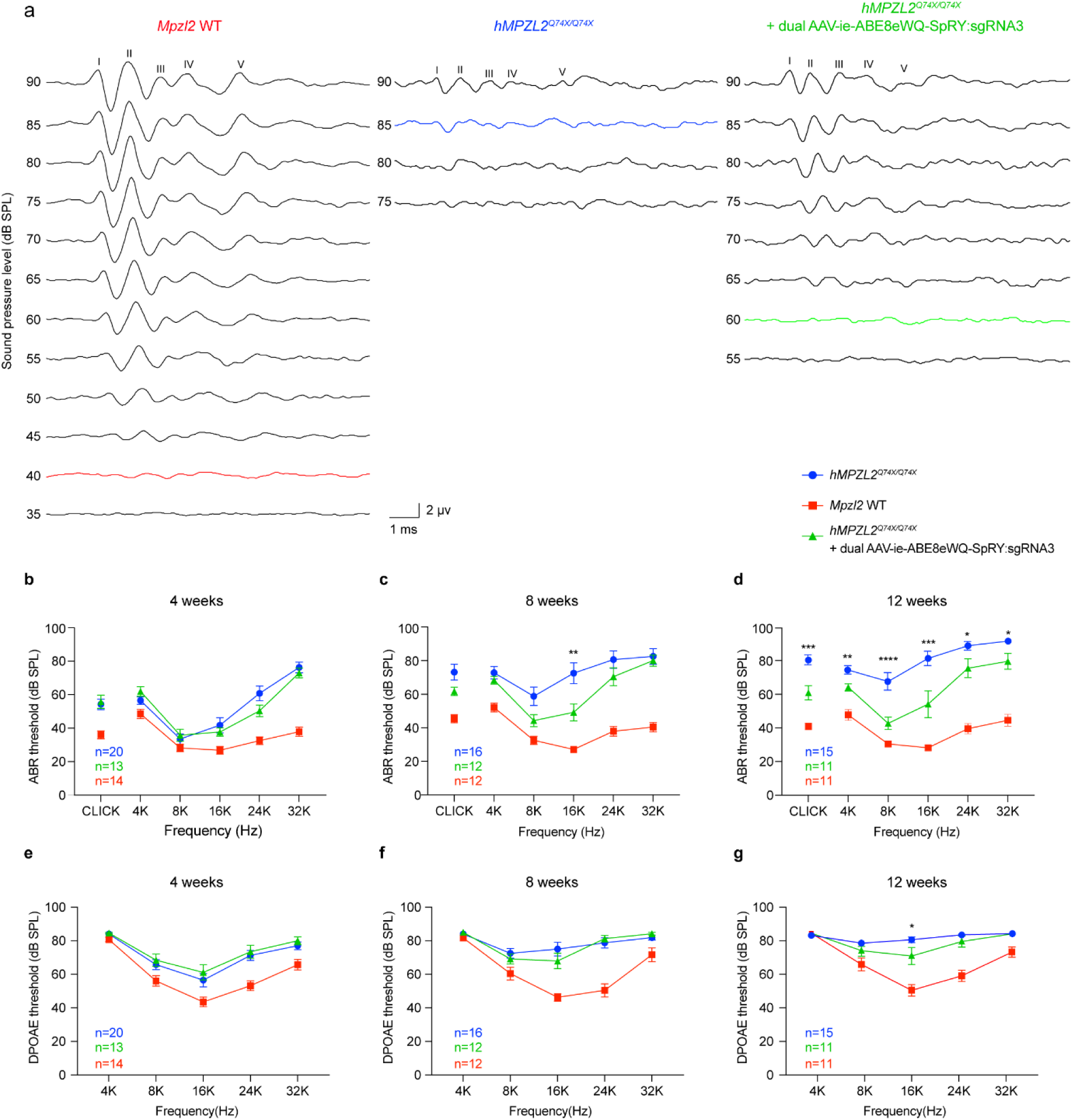
*In vivo* base editing of *MPZL2* restored hearing in *hMPZL2*^Q74X/Q74X^ mice. (a) Representative ABR waveforms from three groups of mice (*Mpzl2*^WT^, *hMPZL2*^Q74X/Q74X^, and treated *hMPZL2*^Q74X/Q74X^) i n response to click stimuli recorded at 12 weeks post-injection. The bold line in each group represents the threshold. (b-d) Comparison of ABR thresholds among *Mpzl2*^WT^, *hMPZL2*^Q74X/Q74X^, and treated *hMPZL2*^Q74X/Q74X^ mice at 4 weeks (b), 8 weeks (c), and 12 weeks (d). (e-g) Comparison of DPOAE thresholds among *Mpzl2*^WT^, *hMPZL2*^Q74X/Q74X^, and treated *hMPZL2*^Q74X/Q74X^ mice at 4 weeks (e), 8 weeks (f), and 12 weeks (g). Statistical significance was calculated using one-way ANOVA with Bonferroni’s correction for multiple comparisons. Significance levels are indicated as \**P* < 0.05, \*\**P* < 0.01, \*\*\**P* < 0.001, and \*\*\*\**P* < 0.0001.

### In vivo correction of the *MPZL2* c.220C>T mutation by dual AAV-ie-ABE8eWQ-SpRY:sgRNA3

To efficiently deliver ABE8eWQ-SpRY:sgRNA3 to the cochlea, we used AAV-ie, a variant with superior inner ear tropism and minimal ototoxicity that is known to target cochlear hair cell (HCs) and supporting cell (SCs) more effectively than other AAV serotypes^11,23^. Given the limitation of AAVs’ small packaging capacity (∼4.7 kb), we used a split intein-mediated AAV system to generate the dual vector of ABE8eWQ-SpRY:sgRNA3. The ABE was split into the amino-terminal (N) and carboxy-terminal (C) halves at amino acid residue 574 of SpRY (**Fig.4a**). Before administering the dual AAV system *in vivo*, we first confirmed that the dual vector encoding each half of ABE8eWQ-SpRY:sgRNA3 had sufficient editing activity in HEK293T-*MPZL2* c.220C>T mutant clonal cells (**Fig.4b**). Approximately 50% on-target editing was achieved, with no bystander edits, which was comparable to that of the single ABE8eWQ-SpRY plasmid. We proceeded to package and deliver each individual AAV vector in AAV-ie capsids. A total of 2 µL dual AAV (1:1 mixed) at a titer of 5.0 × 10^13^ vg/mL was injected into the inner ear of neonatal *hMPZL2*^Q74X/Q74X^ mice at P1-2 via the round window membrane (**Fig.4c**) as previously described^11^.

Fourteen days after injection, the organ of Corti was dissected from the cochleae of treated *hMPZL2*^Q74X/Q74X^ mice. We sequenced the DNA from the cochleae of treated *hMPZL2*^Q74X/Q74X^ mice and tested the base editing efficiency at the *MPZL2* c.220C>T locus in the organ of Corti of eight treated mice. Our results showed that the on-target editing efficiency of treated *hMPZL2*^Q74X/Q74X^ mice was 2.04% ± 0.44% (range: 1.28%–3.66%) (**Fig.4d**), and the bystander editing efficiencies of A0, A1, A2, and A9 were significantly low (**Fig.4e**). We further used the Cas-OFFinder software to search for potential off-target sites of SpRY in mice and implemented high-throughput sequencing to analyze six potential off-target sites. In three treated mice aged 16 days, no obvious off-target editing was detected at the six predicted off-target sites compared with untreated mice (**Fig.4f**). These results indicated that dual AAV-ie-ABE8eWQ-SpRY:sgRNA3 performed with relatively high precision without detectable bystander edits and with clinically relevant editing efficiency.

### In vivo base editing of *MPZL2* restored the hearing of *hMPZL2*^Q74X/Q74X^ mice

To investigate the *in vivo* base editing efficacy on auditory function, ABR and DPOAE thresholds were recorded at 4 weeks, 8 weeks, and 12 weeks post-injection. Representative click ABR waveforms at 12 weeks from *Mpzl2*^WT^, untreated *hMPZL2*^Q74X/Q74X^, and treated *hMPZL2*^Q74X/Q74X^ mice are shown in **Fig.5a.** At 4 weeks, treated *hMPZL2*^Q74X/Q74X^ mice exhibited slightly lower ABR thresholds compared to untreated mice, showing reductions of 4.1 dB, 10.4 dB, and 3.2 dB at 16 kHz, 24 kHz, and 32 kHz, respectively, although these reductions were not statistically significant (**Fig.5b**). We continued to assess hearing function over time and observed more pronounced improvements at 8 weeks and 12 weeks post-injection. At 8 weeks, the ABR thresholds in the treated ears improved by 2.5 dB to 23.3 dB across all frequencies (**Fig.5c**). Specifically, a significant difference in hearing thresholds at 16 kHz was observed between the treated ears of *hMPZL2*^Q74X/Q74X^ mice and untreated *hMPZL2*^Q74X/Q74X^ mice (49.2 ± 5.0 dB vs. 72.5 ± 6.1 dB, *P* < 0.01) (**Fig.5c**). By 12 weeks, the treated ears showed greater recovery compared to untreated mice, with threshold improvements from 12.2 dB to 28.6 dB across all frequencies, showing significant differences (**Fig.5d**). The ABR threshold at 8 kHz in treated *hMPZL2*^Q74X/Q74X^ mice was comparable to that of *Mpzl2*^WT^ mice (**Fig.5d**). The DPOAE thresholds did not show obvious differences between treated and untreated *hMPZL2*^Q74X/Q74X^ mice at 4 weeks (**Fig.5e**). At 8 weeks, the DPOAE thresholds of the treated ears decreased by 3.3 dB and 7.1 dB at 8 kHz and 16 kHz, respectively (**Fig.5f**). At 12 weeks, the decreases were 6.1 dB, 13.6 dB, and 5.0 dB at 8 kHz, 16 kHz, and 24 kHz, respectively, with a significant difference observed at 16 kHz (*P* < 0.05) (**Fig.5g**).

To confirm the safety of our therapeutic strategy, dual AAV-ie-ABE8eWQ-SpRY:sgRNA3 was injected into the inner ears of *Mpzl2*^WT^ mice. The ABR thresholds of injected *Mpzl2*^WT^ ears showed no deterioration compared to untreated *Mpzl2*^WT^ mice and un-injected (contralateral) *Mpzl2*^WT^ ears at 4 weeks, 8 weeks, or 12 weeks (**Extended Data Fig.8a**). Additionally, no degeneration of OHCs or DCs was observed in the organ of Corti of treated mice (**Extended Data Fig.8b, c**). Thus, *in vivo* base editing with the dual AAV-ie-ABE8eWQ-SpRY-sgRNA3 system demonstrated a favorable safety profile with no observed ototoxicity.

### In vivo base editing of *MPZL2* rescued the MPZL2 expression, inner ear structure, and molecular functions in *hMPZL2*^Q74X/Q74X^ mice

The *hMPZL2*^Q74X/Q74X^ mice treated at P1-2 were euthanized at P28, and RT-PCR and immunofluorescence analyses were performed. Human *MPZL2* mRNA expression was significantly restored post-injection compared to untreated mice (**Fig.6a**). Consequently, MPZL2 expression in the cochlea, particularly in the organ of Corti of the treated mice, was significantly rescued, reaching levels comparable to those observed in *Mpzl2*^WT^ mice (**Extended Data Fig.9, Fig.6b**). These data suggest that normal MPZL2 protein was properly constructed and expressed in the target cells post-injection. Furthermore, at 12 weeks the number of OHCs in middle turn and the number of DCs in the middle and basal turns of treated mice were significantly greater than those in untreated mice **(Fig.6c)**. Similarly, we observed increased survival of OHCs and DCs, as well as preserved organ of Corti, in treated mice compared with untreated mice (**Fig. 6d**). Additionally, treated mice exhibited visible hair bundles, particularly in the middle and basal turns (**Fig.6e**). We subsequently performed RNA-seq to identify alterations in gene expression profiles in *hMPZL2*^Q74X/Q74X^ mice treated with dual AAV-ie-ABE8eWQ-SpRY:sgRNA3. Principal Component Analysis of the RNA-seq data revealed a distinct clustering pattern of gene expression between treated and untreated mice (**Fig.6f**). A total of 5,773 DEGs (3,693 upregulated and 2,080 downregulated) were identified between treated and untreated mice (**Extended Data Fig.10a**). GO analysis of the DEGs revealed significant enrichment in biological processes related to cell adhesion, ECM organization, and immune system process (**Fig.6g**). Through a filtering process, we pinpointed the upregulated DEGs in treated *hMPZL2*^Q74X/Q74X^ mice relative to untreated *hMPZL2*^Q74X/Q74X^ mice, identifying 80 transcripts that were also significantly downregulated in untreated *hMPZL2*^Q74X/Q74X^ mice compared with the *Mpzl2*^WT^ mice (**Fig.6h**). These transcripts were notably enriched in biological processes related to cell adhesion and ECM organization (**Extended Data Fig. 10b**). Specifically, we identified 14 transcripts associated with cell adhesion and ECM organization, and gene expression profiles in treated *hMPZL2*^Q74X/Q74X^ mice were restored to the levels observed in *Mpzl2*^WT^ mice, in contrast to the downregulated expression in untreated *hMPZL2*^Q74X/Q74X^ mice (**Fig.6i**). Collectively, these findings demonstrate that dual AAV-ie-ABE8eWQ-SpRY:sgRNA3 treatment not only restores *MPZL2* mRNA and protein expression levels, but also rescues the altered cellular pathways underlying the c.220C>T mutation, resulting in significant improvements in both hearing function and inner ear structural integrity.

**Figure 6.**
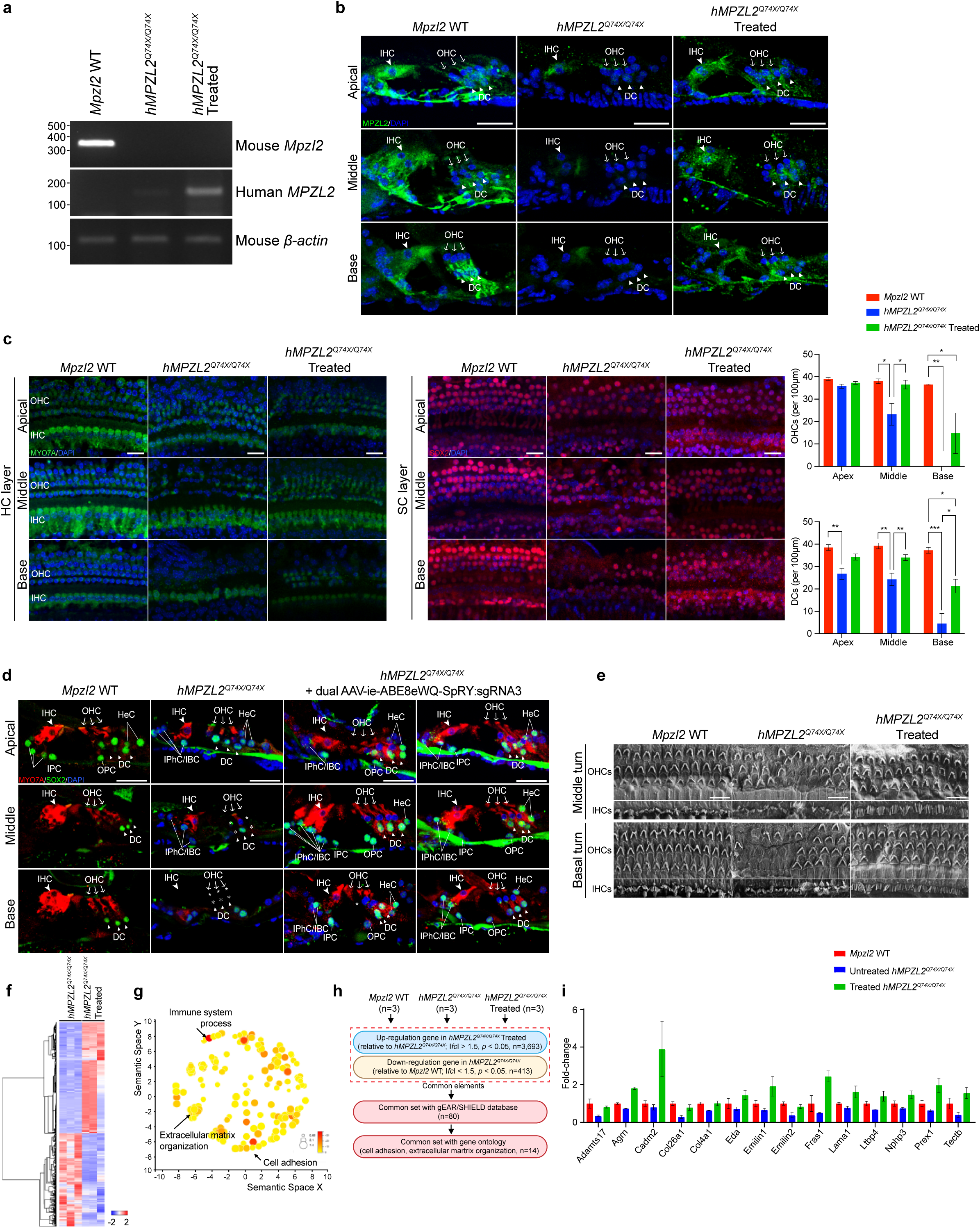
*In vivo* base editing of *MPZL2* rescued MPZL2 expression, inner ear structure, and molecular functions in *hMPZL2*^Q74X/Q74X^ mice. (a) RT-PCR showed enhanced human *MPZL2* gene expression in cochlear membranous tissues at P28 following dual AAV-ie-ABE8eWQ-SpRY:sgRNA3 injection at P2. (b) Representative high-magnification confocal images of cryosection immunofluorescence showing MPZL2 spatial expression in the organ of Corti at P28 following dual AAV-ie-ABE8eWQ-SpRY:sgRNA3 injection at P2. Tissues were immunolabeled with an anti-MPZL2 antibody (green). Scale bar: 25 μm. (c) Representative whole-mount images of the hair cell layer and supporting cell layer from *Mpzl2*^WT^, untreated *hMPZL2*^Q74X/Q74X^, and treated *hMPZL2*^Q74X/Q74X^ mice at 12 weeks of age, depicting the apical, middle, and basal turns, along with the corresponding quantification of surviving OHCs and DCs (the number of animals in each group: n = 4) immunolabeled with anti-Myosin VIIa (HCs, green) and anti-Sox2 (SCs, red). Scale bar: 20 μm. Statistical significance was determined using one-way ANOVA with Bonferroni’s correction for multiple comparisons. Significance levels are indicated as \**P* < 0.05, ** *P* < 0.01, \*\*\**P* < 0.001, and \*\*\*\**P* < 0.0001. (d) Representative section images of the organ of Corti at 12 weeks of age in *Mpzl2*^WT^ (n = 1), untreated *hMPZL2*^Q74X/Q74X^ (n = 1), treated *hMPZL2*^Q74X/Q74X^ mice (n = 2) after injection at P2. The sections were immunolabeled with anti-Myosin VIIa (HCs, red) and anti-Sox2 (SCs, green) antibodies. Scale bar: 25 μm. (e) Scanning electron microscope images of hair bundle morphology from *Mpzl2*^WT^, *hMPZL2*^Q74X/Q74X^, and treated *hMPZL2*^Q74X/Q74X^ mice in the middle and basal turns of the cochlea. Scale bar: 10 μm. (f) Heatmap analysis of DEGs between untreated *hMPZL2*^Q74X/Q74X^ mice (n = 3) and treated *hMPZL2*^Q74X/Q74X^ mice (n = 3) (upregulation, red; downregulation, blue). (g) Revigo visualization of the biological processes (BP) terms associated with cell adhesion, ECM organization, and immune system processes. The size of each circle indicates the relative number of genes in the respective gene set, while the color indicates the significance. (h) Schematic diagram illustrating the RNA-sequencing analysis conducted between *Mpzl2*^WT^ (n = 3), untreated *hMPZL2*^Q74X/Q74X^ (n = 3), and treated *hMPZL2*^Q74X/Q74X^ mice (n = 3), identifying 14 genes of interest associated with cell adhesion and ECM organization. (i) Comparative analysis of the expression levels of the 14 genes of interest related to cell adhesion and ECM organization across the three groups.

## Discussion

Recessive mutations are the most frequent cause of hereditary deafness^9^. Recessive mutations can be treated with AAV-mediated gene replacement, which delivers wild-type cDNA into inner ear cells to correct the underlying genetic defects^24^. We recently used the AAV-ie capsid to deliver the mouse *Mpzl2* cDNA controlled by a ubiquitous promoter into the inner ear of *Mpzl2* knock-out mice and rescued their hearing^11^. However, because the MPZL2 protein is expressed across multiple cell types within the inner ear, and because there is currently a lack of AAV serotypes and specific promoters that can precisely target these cells, gene replacement therapy may result in both overexpression of MPZL2 in targeted cells and ectopic expression in non-targeted cells, potentially leading to cytotoxic effects^25,26^. In addition, it is known that AAV-mediated gene therapies often fail to sustain their therapeutic effects over extended periods^27,28^.

An alternative treatment strategy for hereditary deafness is gene editing, which corrects the mutated DNA bases and addresses the genetic cause of hearing loss at its “root” rather than replacing the mutant gene^29^. Previously, *in vivo* CRISPR nuclease-based gene editing was used to disrupt the reading frame of dominant-negative alleles in dominant deafness mouse models^30–32^ and to restore the nonhomologous end joining-mediated frame in the *Pcdh15* mouse model of DFNB23^33^. However, CRISPR-based disruption of the disease locus is ineffective for ameliorating recessive alleles. Moreover, CRISPR-driven double-strand breaks lead to unintended consequences such as large DNA deletions, chromosomal depletion, and p53-driven programmed cell death^34,35^. To bypass this issue, base editors offer a one-time therapeutic strategy to permanently correct pathogenic mutations without generating double-strand breaks ^36,37^. However, a significant limitation of base editing gene therapy is the need to design distinct sgRNA systems for different mutation sites within the same gene, which demands substantial human and financial resources. In this regard, mutational hotspots or founder mutations present attractive targets for base editors. In DFNB111 patients, the predominant *MPZL2* c.220C>T mutation enables universal application of the dual AAV-ie-ABE8eWQ-SpRY:sgRNA3 system without redesign.

Because inner ear HCs lack regenerative ability in mammals, their degeneration leads to permanent hearing loss. Congenital deafness caused by gene mutations induces disrupted inner ear structural integrity at birth or even during embryonic development, and in such cases gene transfer early in the prenatal period to correct genetic defects and prevent irreversible degeneration is an effective treatment strategy^3^. However, this strategy is complex and carries significant risks, precluding progress in preclinical research on gene therapies for congenital deafness. Preclinical research and clinical translation have advanced more rapidly for genes like *OTOF*, where the mutations do not disrupt the structural integrity of the inner ear^38,39^. Additionally, progressive hereditary deafness is particularly favorable for the development of gene therapy because the target cells in the cochlea remain structurally intact and functionally competent, providing a wider therapeutic window for intervention. Our study, along with previous studies, suggest that *Mpzl2* mutants in mice do not interfere with cochlear development or cause the degeneration of normal HCs and SCs at P4^11,40^. In *hMPZL2*^Q74X/Q74X^ mice, hearing loss and the degeneration of OHCs and DCs both progress from P28 onward, providing a sufficient time window between the onset of hearing loss, the genetic diagnosis, and treatment^41^. Similarly, children with DFNB111 often experience milder forms of hearing loss during their teenage years, with some even passing newborn hearing screenings^12^. Notably, the extended therapeutic window for late-onset progressive deafness might reduce the risk of gene editing delivery vectors dispersing into the brain parenchyma, which is mitigated by the progressive occlusion of communication between the cochlear perilymph and cerebrospinal fluid in the human cochlea^42^.

Key factors – including advanced editing efficacy, safety, and optimized vector tropism – are pivotal in correcting mutant alleles and restoring auditory function. For ABEs, a wild-type tRNA-specific adenosine deaminase (TadA) from *Escherichia coli* was engineered to operate on DNA instead of RNA, and ABEmax with TadA7.10 and ABE8e with TadA-8e variants were developed to have higher base editing activities. However, both ABEmax and ABE8e also have unwanted C-to-T conversion activities at a preferred motif (TC*N) and also have transcriptome-wide sgRNA-independent off-target effects^43^. To address these issues, we previously developed a novel ABE variant (ABE8eWQ) by introducing two variants (V106W and D108Q) into ABE8e, which resulted in significantly reduced bystander effects and RNA off-target activity^43,44^. We found that the c.220C>T mutation could be targeted by ABE8eWQ-SpRY, which demonstrated high editing precision and robust editing efficiency *in vitro*. SpRY has the potential to broadly target the genome, and this is associated with off-target effects at the DNA level^14^. However, ABE8eWQ-SpRY exhibited no significant DNA off-targets at predicted sgRNA-dependent sites, along with undetectable RNA off-targets in the human cells. Using the dual AAV-ie-ABE8eWQ-SpRY:sgRNA3 system, we herein achieved high precision without detectable bystander effects and with relevant editing efficiency *in vivo*. This correlated with the restoration of hearing function, inner ear structures, and the expression of *MPZL2* and related genes. Nevertheless, the ABE-sgRNA therapeutic system still requires further optimization to enhance its editing efficiency *in vivo*, with the goal of restoring higher levels of auditory function and with long term effects.

In summary, we identified an East Asia-specific founder mutation, the homozygous c.220C>T mutation in *MPZL2*, as a promising target for ABE-based gene therapy. The development of humanized mouse models and the successful correction of this mutation using a single PAM-flexible ABE may be a step toward clinical translation of base editing gene therapy for treating hereditary deafness, including most cases of *MPZL2* deafness.

## Acknowledgements

Most analysis of the sequencing data was carried out using the computing server at the Genomic Medicine Institute Research Service Center. This research was supported by grants from the SNU Medicine grant (basic and clinic cooperation research grant No. 800-20230428 to S.-Y.L), by the National Research Foundation (NRF) of Korea (No. 2022R1C1C1003147 to S.-Y.L., No. 2021M3A9H3015389, SRC-NRF2022R1A5A102641311 to S.B.), by the SNUH Kun-hee Lee Child Cancer & Rare Disease Project (No. FP-2022-00001-004 to S.-Y.L), and by the Korean Fund for Regenerative Medicine (KFRM) grant No. RS-2024-00332601 to S.B. The study was also supported by the National Natural Science Foundation of China (grants 82225014, 82171148 to Y.S., grant 82301318 to B.Z., and grant 82301332 to H.T.), the National Key R&D Program of China (grant 2020YFA0908201 to Y.S., and grant 2023YFA0915004 to H.T. and B.Z.), the Science and Technology Commission of Shanghai Municipality (grants 21S11905100 and 23J31900100 to Y.S.), the Shanghai Municipal Health Commission (grant 20224Z0003 to Y.S.), the Shanghai Municipal Education Commission (grant 2023ZKZD12 to Y.S.), Fudan University (grant yg2022-23 to Y.S.), and the Chenguang Program of the Shanghai Education Development Foundation and the Shanghai Municipal Education Commission (grant 23CGA08 to B.Z.). We wish to thank Aavatar Therapeutics (Seunghee Cho, Sangho Park, Dongwoo Song, and Beomseok Choi) for their invaluable technical support during this research.

## Author contributions

Y.S., S.-Y.L., and S.B. jointly conceived and supervised the project. S-Y.L., and S.H.J. performed the sequencing and bioinformatics analysis. S.W.H., H.K., W.H.C., H.S., B.Z., D.M., H.G., Y.Z., H.T., L.C., and X.C. performed the in vitro experiments. L.J., S.J., M.G.K., H.B.C., S-Y.L., Z.W., Y.Z., Z.Z., Y.X., and L.G. performed the in vivo experiments and analyzed the data. Y.S., S.-Y.L., S.B., S.W.H., and L.J. wrote the manuscript. Y.S., S.-Y.L., S.B., S.W.H., L.J., Z.W., A.C., and B.Z. reviewed and revised the manuscript. All authors read and approved the final manuscript.

## Additional information

Supplementary Information accompanying this paper is available at http://

## Materials and Methods

### Human genetics

In this study, we used a retrospective research design and focused on pediatric participants attending the Hereditary Hearing Loss Clinic within the otorhinolaryngology departments of two institutions. The demographic data and audiological phenotypes were retrieved from the electronic medical records. All procedures were approved by the Institutional Review Board of Seoul National University Hospital (no. IRB-H-0905-041-281 and IRB-H-2202-045-1298) and by the Eye, Ear, Nose and Throat Hospital affiliated with Fudan University Review Board of the Office of Research Compliance through protocol 2020122-1 and 2024073. We obtained written informed consent from either the parents of the children or from the participants themselves.

### Whole-exome sequencing and Multiplex ligation-dependent probe amplification

Genomic DNA was extracted from peripheral blood samples using a Chemagic 360 instrument (Qiagen, Venlo, Netherlands) or a DNeasy 96 Blood & Tissue Kit (Qiagen, Germany). We first applied whole-exome sequencing to sequence the exonic regions. The target regions were captured using SureSelectXT Human All Exon V5 (Agilent Technologies, Santa Clara, CA, USA) or CapTruth Human Exome V2.0 (Medical Laboratory of Nantong ZhongKe Co., Ltd., Nantong, China). A library was prepared following the manufacturer’s instructions and was paired-end sequenced using a NovaSeq 6000 sequencing system (Illumina, San Diego, CA, USA) or DNBSEQ-T7 platform (MGI, Shenzhen, China) with an average depth of coverage of 100×. Sequencing reads were aligned to the human reference genome (GRCh38) and processed according to the Genome Analysis Toolkit best-practice pipeline for calling single nucleotide variants and short insertions/deletions (indels)^45^. The ANNOVAR program was used for variant annotation using the RefSeq gene set and Genome Aggregation Database (gnomAD)^46,47^. Rare non-silent variants were selected as candidates, including nonsynonymous single nucleotide variants, coding indels, and splicing variants. We also used the Korean Reference Genome Database (KRGDB) and KOVA (Korean Variant Archive; Korean population database) databases for further filtration of ethnic-specific variants^48,49^. Additionally, the ClinVar and HGMD databases were screened to check whether candidate variants had been previously identified in other patients^50,51^. For individuals exhibiting non-syndromic, symmetric, mild-to-moderate SNHL, we evaluated copy number variations using the SALSA Multiplex Ligation-dependent Probe Amplification Probemix P461-B1 STRC-CATSPER2-OTOA (MRC-Holland, Amsterdam, the Netherlands)^52^.

### Whole-genome sequencing

Patients who remained undiagnosed after whole-exome sequencing were subjected to whole-genome sequencing. DNA libraries were prepared using TruSeq DNA PCR-Free Library Prep Kits (Illumina) or AngTruth-seq EZ DNA Library Preparation Module (for MGI) v2 (Medical Laboratory of Nantong ZhongKe Co. Ltd., Nantong, China) and sequenced on an Illumina NovaSeq6000 or DNBSEQ-T7 platform with an average depth of coverage of 30×. The obtained genome sequences were aligned to the human reference genome (GRCh38) using the BWA-MEM algorithm. PCR duplicates were removed using SAMBLASTER^53^. The initial mutation calling for base substitutions and short indels was performed using HaplotypeCaller and Strelka2, respectively.^54^ Structural variations were identified using Delly^55^. We classified candidate variants according to the American College of Medical Genetics and Genomics and Association for Molecular Pathology guidelines for hearing loss^56^.

### Humanized mice

C57BL/6N and ICR mice were purchased from Orientbio Inc. (Sungnam, Republic of Korea) and were housed under specific pathogen-free conditions in individually ventilated cages. The temperature and humidity of the breeding environment were maintained at 22 ± 1 °C and 50%, respectively, with a 12 hour light:dark cycle. All feed and individually ventilated cages were sterilized, and all air conditioners had filters. The protocols for the generation of *hMPZL2*^Q74X^ and *hMPZL2*^WT^ mice were performed according to the Korean Food and Drug Administration (KFDA) guidelines and were reviewed and approved by the Institutional Animal Care and Use Committees (IACUC; MS-2022-01, MS-2023-01) at the GEM Center of Macrogen (Seoul, Republic of Korea) and at Seoul National University Hospital.

### Humanized mouse sgRNA

sgRNA searches and potential off-target analyses were performed using the CRISPR design tools CHOPCHOP (https://chopchop.cbu.uib.no)^57^ and CRISPOR (https://crispor.gi.ucsc.edu)^58^. To generate the *hMPZL2*^Q74X^ mouse, the sgRNA was searched based on the NCBI database (NC_000075.7, NM_007962.4) within the partial regions of exon 1 and intron 1, including the 5’ UTR-ATG region. For the *hMPZL2*^WT^ mouse, the search was based on human *MPZL2* (*hMPZL2*) transcript NM_005797.4, within the regions of Exon 2 and 3. The sequences of the sgRNAs used are listed in **Supplementary Table 4.**

### Humanized mouse targeting donor

The *hMPZL2*^Q74X^ targeting donor was designed based on the target site (GRCm39; chr9: 44,954,049) and incorporated approximately 600-bp homology arms on both sides of the *hMPZL2*^Q74X^ expression cassette. The 5’ and 3’ homology arms included sequences from the upstream-5′UTR and from intron 1 of the mouse *Mpzl2* gene, respectively. The designed donor sequences were synthesized in vector DNA form (Invitrogen, Waltham, MA, USA). For the *hMPZL2*^WT^ mouse, single-stranded DNA (ssDNA) was used for the donor. The ssDNA was designed with the *hMPZL2* mutation site (c.220T>C:p.X74Q) at the center and 60bp homology arms on both sides and was synthesized.

### Humanized mouse embryo collection

To induce superovulation, 7.5 IU each of pregnant mare serum gonadotropin (PMSG; Dsmbio, Uiwang, Republic of Korea) and human chorionic gonadotropin (hCG; Dsmbio) were intraperitoneally injected for 5–8 weeks in female mice at 48-hour intervals. Subsequently, zygotes were obtained from the oviducts of these female mice after mating with males to produce *hMPZL2*^Q74X^ mice. To generate *hMPZL2*^WT^ mice, superovulation was induced in the same manner, and then unfertilized embryos were collected from female mice that had not mated with male mice. The collected embryos were in vitro fertilized using sperm from *hMPZL2^Q74X^* homozygous mice to ensure the development of zygotes. Previously reported detailed procedures for in vitro fertilization were referenced^59^. The zygotes were washed in Quinn’s Advantage Medium with HEPES (ART-1024, Cooper Surgical, Trumbull, CT, USA) and then transferred to droplets of KSOM medium (MR-121-D, Sigma-Aldrich, St. Louis and Burlington, MA, USA) in a CO_2_ incubator at 37°C.

### Humanized mouse CRISPR mixture and microinjection

sgRNA was synthesized using the GeneArt Precision sgRNA Synthesis Kit (A29377, Thermo Fisher Scientific, Waltham, MA, USA) according to the manufacturer’s instructions. Microinjection was performed by injecting a mixture composed of Cas9 protein (Macrogen), purified sgRNAs, and linearized *hMPZL2*^Q74X^ vector or *hMPZL2*^WT^ ssDNA into the pronucleus of 1-cell zygotes in Quinn’s Advantage Medium with HEPES at concentrations of 20 ng/µl, 25 ng/µl, and 20ng/µL, respectively. Microinjected zygotes were incubated at 37°C for 2 hours and were transplanted by surgical methods into the oviducts of pseudopregnant recipient ICR mice at the 1-cell or 2-cell stages.

### Humanized mouse genotyping

Genomic DNA for genotyping was extracted from the tails of 2-week-old mice using an Axen Total DNA mini kit (MG-P005-50, Macrogen). The PCR assay was conducted using the EF-Taq DNA Polymerase (SEF16-R250, Solgent, Daejeon, Republic of Korea) and Axen High-Q Taq DNA Polymerase (MG-E-003-250, Macrogen). The PCR was performed in a total volume of 20 µL consisting of 2 µL of 10× reaction buffer, 4 µL of 5× GC enhancer, 0.4 µL of 10 mM dNTP mixture, 2 µL each of 10 µM forward and reverse primers, about 200 ng of extracted genomic DNA, and PCR-grade water to make up the final volume. The PCR cycling conditions were an initial denaturation at 95°C for 5 minutes followed by 35 cycles at 95°C for 10 s, 56–60°C for 20 s, and 72°C for 30 s/kb. For genotyping *hMPZL2*^WT^ mice, PCR products using the *hMPZL2*^Q74X^ allele as a template were added to create heteroduplexes through re-annealing, followed by treatment with T7 endonuclease 1 (M0302S, NEB, Ipswich, MA, USA). For *hMPZL2*^WT^ mice, PCR cloning (E1203S, NEB) was additionally performed for allele separation, and Sanger sequencing was performed for sequence verification of the two types of humanized mice. The sequences of the primers used for genotyping are listed in **Supplementary Table 5**.

### HEK293-*MPZL2* c.220C>T clonal cells

HEK293T cells were purchased from the ATCC and maintained in a strictly mycoplasma-free environment and at low passage numbers. The HEK293T-MPZL2 c.220C>T stable cell line was established by cloning a 201-bp disease-associated gene fragment containing the C·G-to-T·A mutation from the ClinVar database (https://www.ncbi.nlm.nih.gov/clinvar/) by assembling the fragments into a modified lentivector from lentiCRISPR v2 (#52961), thus yielding the lentivector plasmid Lenti MPZL2-P2A-puro. HEK293T cells were seeded into 6-well plates (Corning) at approximately 85% confluency per well and were co-transfected with 2.25 μg of the lentivector plasmid, 2.25 μg psPAX2 (#12260, encoding the viral packaging proteins), and 1.5 μg pMD2.G (#12259, encoding the VSV-G envelope protein) using Lipofectamine 3000 (L3000015, Thermo Fisher Scientific) following the manufacturer’s instructions. Virus-containing supernatant was collected after 48 h of transfection and centrifuged at 2,000 × *g* for 10 minutes to remove cell debris and filtered through a 0.45-μm polyvinylidene difluoride (PVDF) filter (IPVH00010, Millipore). A total of 150 μL filtered virus-containing supernatant was added to HEK293T cells at approximately 40–50% confluency cultured in 6-well plates. After 24 h of transduction with lentivirus, the cells were split into the wells of a new plate supplemented with puromycin (2.5 μg/ml). After 72 h of puromycin selection, a stable cell line with the *hMPZL2* c.220C>T mutation was successfully established. To ensure single-copy integration, cells with the fewest surviving colonies were collected and subsequently expanded for further transfection procedures.

### Cell culture

Cultured cells used in this study, including HEK293-*MPZL2* c.220C>T and patient-derived cell lines, were maintained in a humidified atmosphere containing 5% CO_2_ at 37°C. All culture media consisted of DMEM supplemented with heat-inactivated FBS (10% for HEK293 cells and 20% for patient-derived fibroblasts), 100 units/mL penicillin/streptomycin, and 2 mM L-glutamine. Cells were routinely tested for mycoplasma contamination.

### ABE expression plasmid construction

The ABE8e (no.138489, Addgene, Watertown, MA) and NG-ABE8e (no.138491, Addgene) used in this study were separately obtained from previously reported plasmids available from Addgene. Other ABE variant expression plasmids were constructed by replacing the deaminase and Cas-nuclease domains using a Gibson assembly of linearized destination vector and PCR amplicons with Gibson-compatible ends for each assembly junction. PCR was performed using KOD-Multi & Epi (KME-101, TOYOBO).

### sgRNA expression plasmid construction

The sgRNA expression plasmids for the various Cas effectors were generated using custom oligonucleotides and restriction enzyme-based classical cloning methods based on pRG2 (no. 104174, Addgene). Briefly, for each guide RNA sequence, a pair of complementary oligos with 4-bp overhangs were annealed and inserted via a cut-ligation reaction with BsaI and T4 DNA ligase in the pRG2-hU6 plasmid. All sgRNA sequences were designed with a G preceding 19-bp or 23-bp spacer targets for transcription initiation from the Polymerase III promoter. To produce a vector expressing sgRNA for the other SpCas9 ortholog, eNm2-C, an sgRNA scaffold was synthesized from Macrogen and integrated with the pRG2-hU6 plasmid. The gRNA was inserted again into the plasmid, where the original sgRNA scaffold was swapped with the scaffold tailored for eNme2-C. The sgRNAs were inserted into the PSK-mU6-sgRNA plasmid between two Bbs1 restriction sites. All plasmids used in this study were purified using Nucleobond Xtra Midi EF (MN, 740420.5) for Midi-prep or Exfection plasmid LE (111-102, GeneAll) for Mini-prep. The sequences of the sgRNAs are listed in **Supplementary Table 4**.

### Targeted deep sequencing

To evaluate the on/off-target editing efficiency, genomic DNA was amplified by a three-step nested PCR using KOD-Multi & Epi (KME-101, TOYOBO). PCR 1-2 steps were performed with primers specifically targeting the genomic region of interest. The total PCR cycles were kept to a minimum to avoid PCR bias. The primers used for PCR step 2 included the minimal adapters for adding barcodes in the following PCR step 3. Finally, the barcoded PCR products were pooled, purified using the Expin PCR SV mini kit (103-102, GeneAll), and sequenced using an Illumina Miniseq according to the manufacturer’s instructions. Sequencing results were analyzed using a BE-Analyzer (http://www.rgenome.net/be-analyzer/)^60^.

### RNA off-target analysis

HEK293T cells were seeded in a 24-well plate, and the next day they were transfected with 390 ng ABE-encoding plasmid and 125 ng sgRNA using JetOPTIMUS® (Polyplus) according the manufacturer’s protocol. Briefly, the plasmids were mixed with JetOptimus reagent in JetOptimus buffer, then the mixture was incubated for 10 min and added into each well. For RNA extraction, cells were harvested by treating with TRIzol reagent 24 h after transfection. To synthesize cDNA, reverse transcription was performed using ReverTra Ace-α-(FSK-101, TOYOBO) according to the manufacturer’s instructions. The target region was amplified using KOD-Multi & Epi (KME-101, TOYOBO), and the PCR products were analyzed using an Illumina MiniSeq instrument. To obtain the percentage of adenosines edited to inosines, the number of adenosines converted to guanosines was divided by the total number of adenosines in the products. The sequences of the primers are listed in **Supplementary Table 5**.

### AAV plasmid construction

For AAV production, we used the split intein-mediated ABE constructs designed in a previous study^61^, with some modifications as follows. Dual ABE constructs were amended sequentially in two steps each. The N-terminal vector was first modified by installing an additional A61R mutation (GCG to AGG) in the 5′ half of the Cas9 coding sequence (2–1368 aa) for SpRY using the QuickChange mutagenesis method^62^. The TadA8eWQ domain and linkers were then subcloned into the N-terminal vector between the NotI and BgIII sites. The C-terminal vector was modified to encode the final selected sgRNA3 (NAN PAM) from the in vitro screening and to contain the remainder of the SpRY mutations. A pair of recombinant vectors contained either inverted terminal repeat (ITR)-CMV promoter-nucleic localization signal (NLS)-ecTadA8eWQ-SpRY(N)-Npu(N)-bGHpA-ITR or ITR-CMV promoter-Npu(C)-SpRY(C)-NLS-1xHA-bGHpA;U6-sgRNA-ITR. The dual viral plasmids for AAV packaging were purified using Nucleobond Xtra Midi EF (MN, 740420.5) for Midi-prep, and the dual viral plasmids were separately packaged into two AAV-ie viruses (PackGene Biotech).

### Round window injection into the inner ear of neonatal mice

Neonatal mice of either sex were used for injections, and the mice were randomly assigned to the different experimental groups. All surgical procedures were performed in a clean, dedicated space, and all instruments were thoroughly cleaned with 70% ethanol and autoclaved prior to surgery. P1-2 *hMPZL2*^Q74X/Q74X^ mice were used for AAV-ie-ABE8eWQ injection. Mice were anesthetized by hypothermia on crushed ice, and a skin incision was made behind the ear of the mouse to expose the tympanic ring and the stapedial artery under an operating microscope (OPMI pico, ZEISS). Glass micropipettes (WPI) held by a Nanoliter 2020 Microinjection System (WPI) were inserted through the round window membrane, which allows access to inner-ear cells. The total injection volume was 2 μl per cochlea for round window membrane injection, and the release rate was 7 nL/s under the control of a MICRO2T SMARTouch microinjection controller (WPI). The skin was closed with a 6-0 nylon suture (NB617P, Ailee).

### In vivo gene editing efficiency test

Genomic DNA was purified from the organ of Corti of injected *hMPZL2* c.220C>T mice and untreated *hMPZL2* c.220C>T mice (negative control). Purified DNA was amplified by PCR for each condition using primers for the genomic DNA. HTS amplicon libraries containing an adaptor sequence (forward, 5′-ACA CTC TTT CCC TAC ACG ACG CTC TTC CGA TCT-3′; reverse 5′-ACT GGA GTT CAG ACG TGT GCT CTT CCG ATC T −3′) at the 5′ end were prepared by PCR using KOD-Plus-Neo DNA Polymerase (KOD-201, Toyobo). The aforementioned products were processed through another round of PCR amplification with distinct barcode sequences added to the primers. The resulting libraries were pooled and subjected to 150 bp paired-end sequencing on an Illumina HiSeq platform. The A-to-G conversions in the HTS data were analyzed with a BE-Analyzer^60^.

### Audiometric tests

Mice were anesthetized via intraperitoneal injection with a mixture of dexmedetomidine hydrochloride (200 mL/kg) and Zoletil (35 mg/kg). ABR and DPOAE thresholds were measured using a BioSigRZ system (Tucker-Davis Technologies, Alachua, FL, USA) in a soundproof chamber. Anesthetized animals were first recorded to collect ABR, and then moved to a second setup to record DPOAE.

### ABR recordings and analysis

ABR signals were collected via subdermal needle electrodes placed at the mastoid portion of the skull, the vertex of the skull, and the dorsal rump as the recording electrode, reference electrode, and ground electrode, respectively. The ABR signals were evoked and filtered through a 300 Hz to 3 kHz passband and averaged at 512 repeats of each sound pressure level (SPL). Click stimuli (90 to 10 dB, 2.5 ms, 21 pps rate) and tone-burst stimuli (4, 8, 16, 24 and 32 kHz, 2.5 ms, 21 pps rate) were applied from 90 to 10 dB SPL in 10 dB steps. The ABR threshold was defined as the lowest sound level at which any waveform peak could be observed.

### DPOAE recordings and analysis

The f1 and f2 primary tones were generated at a frequency ratio of 1.2 with f2 levels at 4, 8, 16, 24, and 32 kHz and L1 – L2 = 10 dB SPL. The f2 levels were swept in 5 dB steps from 80 dB to 20 dB SPL. The DPOAE threshold was defined from the average spectra at the f2 levels that produced a DPOAE with a magnitude 5 dB SPL above the noise floor.

### Histology

Histological and immunofluorescence analyses were conducted on cochleae from 12–15-week-old mice. After anesthetization via intraperitoneal injection, the mice were perfused with 4% paraformaldehyde (BPP-9004-001L, T&I) and the cochleae were dissected from the temporal bone. For histology, the cochleae were decalcified and embedded in paraffin blocks that were then sectioned at a thickness of 4 μm using a rotary microtome. The tissue sections were mounted on glass slides, deparaffinized in Histo-clear II (HS-202, National Diagnostics) for 10 minutes, washed in 100%, 95%, 90%, and 70% ethanol, and finally washed for 10 minutes in tap water. Hematoxylin and eosin staining was performed, with the slides being incubated in hematoxylin for 2 minutes and then stained with eosin for 20 seconds. The slides were subsequently mounted using a mounting medium and analyzed for cytoarchitecture evaluation at 4×, 20×, and 100× magnification using an ECLIPSE Ci microscope (Nikon).

### Immunofluorescence

For Immunofluorescence assays of 12–15-week-old cochlea, tissue samples were post-fixed with 4% paraformaldehyde overnight at 4°C. The cochleae were rinsed with PBS and decalcified using 10% ethylene diamine tetraacetic acid (EDTA) overnight at room temperature (RT). The decalcified cochleae were cryoprotected with a sucrose gradient in PBS (10%, 20%, and 30% sucrose) for 1 h at RT, then soaked in 30% sucrose overnight at 4°C. The specimens were embedded in Tissue-Tek OCT, and mid-modiolar cryosections were made at 10 µm thickness and mounted on glass slides. The tissue sections were permeabilized with 1% Triton X-100 and blocked with 4% BSA in PBS for 20 minutes at RT and then incubated overnight at 4°C with the following primary antibodies: rabbit anti-MYO7A (1:400 dilution, PA1-936, Invitrogen), mouse anti-SOX2 (1:400 dilution, sc-365823, Santa Cruz), rabbit anti-MPZL2 (1:300 dilution, 11787-1-AP, Proteintech), and mouse anti-TUBB3 (1:300 dilution, 801202, Biolegend). After washing with PBS, the sections were incubated with the corresponding Alexa-conjugated secondary antibodies for 1 h at RT. After rinsing with PBS, specimens were mounted on microscope glass slides using mounting medium with DAPI (AB104139, Abcam). Images were captured using a Zeiss Axioskop light microscope and a Leica TCS SP5 confocal microscope (Leica, Wetzlar, Germany). For whole-mount immunofluorescence staining, the decalcified cochleae were dissected into three pieces in ice-cold PBS. Samples were permeabilized, blocked, and incubated overnight with rabbit anti-MYO7A and mouse anti-SOX2 primary antibodies at 4°C. Appropriate Alexa-conjugated secondary antibodies were applied for 1 h at RT, and mounting medium with DAPI was used to mount the specimens. Images were obtained using a Leica TCS SP8 confocal microscope (Leica, Wetzlar, Germany). The numbers of MYO7A^+^ inner hair cells and OHCs and SOX2^+^ DCs were counted in 100 μm cochlear sections across the apical, middle, and basal turns. The number of SGNs in the apical, middle, and basal turns was counted using ImageJ software. For SGN quantification, TUBB3^+^ SGNs were counted per 72,900 μm². SGN density in H&E-stained paraffin sections was determined by counting the number of SGNs per 8,000 μm².

### Scanning electron microscopy

Cochleae were dissected and gently perfused with 2.5% glutaraldehyde (G7651-10ML, Sigma-Aldrich) in PBS through the round and oval windows and into the apex. Samples were fixed in 2.5% glutaraldehyde in PBS overnight at 4°C. After washing with PBS, the cochleae were decalcified and dissected into three turns. The specimens were post-fixed in 1% osmium tetroxide (19172, EMS) in PBS for 2 hours and dehydrated in an ethanol gradient (50%, 60%, 70%, 80%, 90%, and 100% ethanol) at 10-minute intervals. The samples were then rinsed with a mixture of 100% ethanol and hexamethyldisilazane (HMDS, Sigma-Aldrich) in a 1:1 ratio for 30 minutes, followed by drying with HMDS only for 30 minutes. The dried samples were mounted on metal disks and gold sputter-coated (Cressington sputter coater 208HR), and images were obtained at 5.0 kV magnification using an electron microscope (Hitachi S-4700 FESEM).

### Reverse transcription (RT)-PCR and quantitative RT-PCR

Cochlea, kidney, and lung tissues were freshly dissected from P4 wild-type and mutant mice. These tissues were immediately frozen in liquid nitrogen and stored at –80°C until processing. Total RNA was isolated using TRIzol Reagent (Invitrogen) as per the manufacturer’s instructions. cDNA synthesis was performed from 2 μg of total RNA using Accupower RT-pre-mix (Bioneer, Daejeon, Republic of Korea). PCR was then performed using diluted cDNA and 10 pmol of specific primers to assess the mRNA levels. The thermal cycling conditions consisted of an initial denaturation at 95°C for 5 min followed by 30 cycles of 95°C for 30 s, 55°C for 30 s, and 72°C for 1 min. The amplified DNA was visualized using agarose gel electrophoresis with Loading STAR (DYNE Bio, Seongnam, Republic of Korea). For quantitative RT-PCR reactions, 1/20 diluted cDNA was combined with SYBR qPCR master mix as the reporter dye and 10 pmol of primers to detect mRNA expression of specific genes. The thermal cycling conditions were as follows: 95°C for 3 minutes for enzyme activation followed by 40 cycles of 95°C for 10 seconds, 53°C for 15 seconds, and 72°C for 30 seconds. The primer sequences are listed in **Supplementary Table 5**.

### RNA sequencing

RNA sequencing analyses were conducted as previously reported^63^. Briefly, total RNA was extracted from P28 mouse cochleae (three pairs of cochleae from wild-type and mutant mice) utilizing TRIzol (Invitrogen, USA) following the manufacturer’s protocol. RNA sequencing libraries were prepared using a SMARTER stranded total RNA-seq in accordance with the manufacturer’s protocols (Takara, Japan). After confirming the library size (approximately 300–500 bp) and fluorophore-based quantification of each library concentration (Quantus; Promega, USA), the samples were sequenced on a Nextseq 2000 under paired-end conditions (Illumina). After filtering, fastq files were mapped against the reference genome GRCm39 (ensemble) and pseudo-aligned by Kallisto (version 0.51.0)^64^. Transcript counts were imported at the gene level using txiimport^65^, and DEGs were identified using DEseq2^66^. Individual samples were further analyzed to list genes with a fold change greater than 1.5 and a p-value < 0.05. DAVID GO analysis was performed to examine GO terms for biological processes, cellular components, and molecular functions^67^. Representative GO terms were summarized and visualized using REVIGO^68^. DEGs were further compared with the mouse cochlear single-cell RNA-seq database (http://umgear.org) and the SHIELD database (http://shield.hms.harvard.edu) in order to refine the list of genes implicated in specific cellular pathways. The RNA-seq findings were validated by quantitative RT-PCR analysis using specific primers for selected DEGs. The sequences of the primers are listed in **Supplementary Table 5**.

### Statistics

Statistical analyses and visualizations were conducted using R Version 4.2.2. and GraphPad Prism (version 9.0.2). The code for human genetics plots **(Fig. 1 and Extended Data Fig. 1)** is available at https://github.com/SNUH-hEARgeneLab/MPZL2. Differences between two groups were analyzed using an unpaired Student’s t-test, while one-way ANOVA with Bonferroni correction and the Kruskal–Wallis test were conducted for comparisons among more than two groups, as appropriate. All data are presented as the mean ± SEM. Statistical significance is indicated in the figures as follows: ns, not significant; \**P*<0.05; \*\**P*<0.01; \*\*\**P*<0.001; \*\*\*\**P*<0.0001.

## Extended data figure legends

**Extended Data Fig. 1.** Mutational landscape of the *MPZL2* mutations causing DFNB111, and the natural progression of hearing loss across different hearing frequencies. (a) The mutational distribution and prevalences of DFNB111, as documented in both the literature and in our cohort. The Circos plot showing the frequency of *MPZL2* mutations depending on the exons (exon 15) and functional domains. (b) The annual progression of hearing loss in DFNB111 patients across different hearing frequencies was determined through linear regression analyses using auditory function-gene profiles (n = 114).

**Extended Data Fig.2.** Validation of the humanized *MPZL2* c.220C>T knock-in mouse model. (a) PCR assay of the *hMPZL2*^Q74X^ mouse. The integration of the human cassette was confirmed using the Fw1-Rev1 primer pair to identify candidates, and final on-targeting was verified using the Fw2-Rev2 primer pair (M: DNA ladder, WT: wild-type, PC: Q74X donor). (b) Low-magnification section images showing *MPZL2* expression in the cochlear turns of *Mpzl2*^WT^ and *hMPZL2*^Q74X/Q74X^ mice at P4 and P28. In *Mpzl2*^WT^ mice, MPZL2 was clearly observed in the organ of Corti at both P4 and P28. However, no expression was detected in *hMPZL2*^Q74X/Q74X^ mice at either time point. The sections were immunolabeled with MPZL2 (green) and DAPI (blue). Scale bar: 100 μm. (c) High-magnification images of organ of Corti sections and whole-mount assays showed distinct MPZL2 expression patterns between *Mpzl2*^WT^ and *hMPZL2*^Q74X/Q74X^ mice at P4 and P28. Scale bar: 25 μm

**Extended Data Fig.3.** Generation and validation of the humanized *MPZL2*^WT^ mouse model. (a) The *hMPZL2*^WT^ mouse was produced using X74Q ssDNA by in vitro fertilization (IVF) and microinjection methods. The black box and line indicate the position of the sgRNA with PAMs, and the red letter and red arrowheads indicate the start and stop codon and the Cas9 cleavage sites, respectively. (b) Gene mutation assay for *MPZL2*^WT^ mice. After amplifying the PCR product using the Fw3-Rev3 primer pair, the Q74X PCR product was added to form a heteroduplex. A cleavage was then confirmed using a T7 endonuclease 1 (T7E1) assay. “–/+ Q74X” indicates the presence or absence of the Q74X PCR products (Het: *hMPZL2*^Q74X^ heterozygote, Homo: *MPZL2*^Q74X^ homozygote, T7P: T7E1 control). (c) Sequence verification results for humanized WT mice by Sanger sequencing.

**Extended Data Fig.4.** Inner ear structure deficits of *hMPZL2*^Q74X/Q74X^ mice. (a) Microphotographs showing representative images of mid-modiolar sections of the cochlea from an *Mpzl2*^WT^ mouse (a’), an *hMPZL2*^Q74X/WT^ mouse (b’), and two *hMPZL2*^Q74X/Q74X^ mice (c’, d’). The apical, middle, and basal turns are boxed in images (a’–d’). Scale bar: 500 μm. Close-up views of the organ of Corti from an *Mpzl2*^WT^ mouse (e’, i’, m’), an *hMPZL2*^Q74X/WT^ mouse (f’, j’, n’), and two *hMPZL2*^Q74X/Q74X^ mice (g’, h’, k’, l’, o’, p’) are shown. Asterisks (*) indicate loss of outer hair cells (OHCs) and Deiter’s cells (DCs). Rectangles indicate the misalignment and disorganization of OHCs and DCs in the organ of Corti (g’, h’) as well as the collapsed tunnel of Corti (o’, p’). BM: basilar membrane. Scale bar: 200 μm. (b) Images of lateral wall and spiral ganglion neuron region sections of the cochlea from an *Mpzl2*^WT^ mouse, an *hMPZL2*^Q74X/WT^ mouse, and two *hMPZL2*^Q74X/Q74X^ mice are shown. Close-up views (rectangles) of the middle region of the stria vascularis and the center of spiral ganglion neurons. Spiral ganglion neuron, SGN; Stria vascularis, SV; Lateral wall, LW; Spiral ligament, SL; Otic capsule, OC. Scale bar: 20 μm. (c) Quantification of SV thickness in *Mpzl2*^WT^, *hMPZL2*^Q74X/WT^, and *hMPZL2*^Q74X/Q74X^ mice at 12 weeks of age. (d) Quantification of SGN density in *Mpzl2*^WT^, *hMPZL2*^Q74X/WT^, and *hMPZL2*^Q74X/Q74X^ mice at 12 weeks of age. Statistical significance was determined using Kruskal–Wallis tests with Dunn’s multiple comparisons tests. ns, no statistical significance.

**Extended Data Fig.5.** Immunohistochemistry of SGN populations of *Mpzl2*^WT^, *hMPZL2*^Q74X/WT^, and *hMPZL2^Q74X/Q74X^* mice at 12 weeks of age. (a) Representative high-magnification confocal images of SGNs stained with anti-TUBB3 (green) from the apical, middle, and basal turns of the cochleae for each genotype. Scale bar: 25 μm. (b) Quantification of SGN density among *Mpzl2*^WT^, *hMPZL2*^Q74X/WT^, and *hMPZL2*^Q74X/Q74X^ mice at 12 weeks of age. Statistical significance was determined using Kruskal–Wallis with Dunn’s multiple comparisons test. ns, no statistical significance.

**Extended Data Fig.6.** Auditory phenotype of the humanized *MPZL2*^WT^ mice (*hMPZL2*^WT/WT^). (a) Representative click-evoked ABR traces in 4, 8, and 12-week-old *Mpzl2*^WT^ (red) and *hMPZL2*^WT/WT^ (blue) mice. The bold line in each trace indicates the hearing threshold. (b) Average ABR thresholds of *Mpzl2*^WT^ mice (red, n = 6) and *hMPZL2*^WT/WT^ (blue, n = 6) across click and tone-burst frequencies at 4, 8, and 12 weeks of age. Data are presented as the mean ± SEM. Statistical significance was determined using the Mann–Whitney U-test with corrections following Student’s t-test. ns, no statistical significance. (c) Representative section images of the organ of Corti in 12-week-old *hMPZL2*^WT/WT^ mice. Immunolabeled with anti-Myosin VIIa (HCs, red), and anti-Sox2 (SCs, green). Scale bar: 25 μm. (d) Representative whole-mount assay at 12-week-age of *hMPZL2*^WT/WT^ mice. Immunolabeled with anti-Myosin VIIa (HCs, red), and anti-Sox2 (SCs, green). Scale bar: 25 μm.

**Extended Data Fig.7.** Gene Ontology (GO) enrichment analysis. GO enrichment analysis was performed in the categories of Biological Processes (green), Cellular Components (red), and Molecular Functions (blue) when comparing *Mpzl2*^WT^ and *hMPZL2*^Q74X/Q74X^ mice at P28.

**Extended Data Fig.8.** Evaluation of the safety of dual AAV-ie-ABE8eWQ-SpRY:sgRNA3 on hearing function and cochlear structure in *Mpzl2*^WT^ mice. (a) Comparison of ABR thresholds of untreated *Mpzl2*^WT^ mice (n = 5), treated mice (injected ears, n = 5), and treated mice (contralateral un-injected ears, n = 5) at 4 weeks, 8 weeks, and 12 weeks. Data are shown as the mean ± SEM. Statistical significance was determined using Kruskal–Wallis tests with Dunn’s multiple comparisons tests. ns, no significance. (b) Representative images of organ of Corti sections at 12 weeks of age in *Mpzl2*^WT^ mice post-injection immunolabeled with anti-Myosin VIIa (HCs, red) and anti-Sox2 (SCs, green) antibodies. Scale bar: 25 μm. (c) Representative whole-mount assays of 12-week-old *Mpzl2*^WT^ mice post-injection immunolabeled with anti-Myosin VIIa (HCs, red) and anti-Sox2 (SCs, green) antibodies. Scale bar: 25 μm.

**Extended Data Fig.9.** Representative low-magnification confocal images of cryosection immunofluorescence showing MPZL2 spatial expression in the cochlea at P28 following dual AAV-ie-ABE8eWQ-SpRY:sgRNA3 injection at P2. Tissues were immunolabeled with an anti-MPZL2 antibody (green). Rectangles highlight the organ of Corti. Scale bar: 100 μm.

**Extended Data Fig.10.** RNA-sequencing analysis. (a) Volcano plot showing differentially expressed genes (DEGs) between untreated *hMPZL2*^Q74X/Q74X^ mice (n = 3) and treated *hMPZL2*^Q74X/Q74X^ mice (n = 3). The number of upregulated genes (red box) and downregulated genes (blue box) is indicated. (b) Gene Ontology (GO) enrichment analysis of differentially expressed genes in biological processes (BP) terms. The analysis focused on 80 transcripts that were significantly downregulated in untreated *hMPZL2*^Q74X/Q74X^ mice compared to *Mpzl2*^WT^ mice and upregulated in treated *hMPZL2*^Q74X/Q74X^ mice. Red boxes highlight BP terms related to cell adhesion and ECM organization, with associated transcript information described.

