## Extended Data Figs for "PAM-flexible adenine base editing rescues hearing loss in a humanized *MPZL2* mouse model harboring an East Asian founder mutation"

### Extended Data Fig.1

**a**

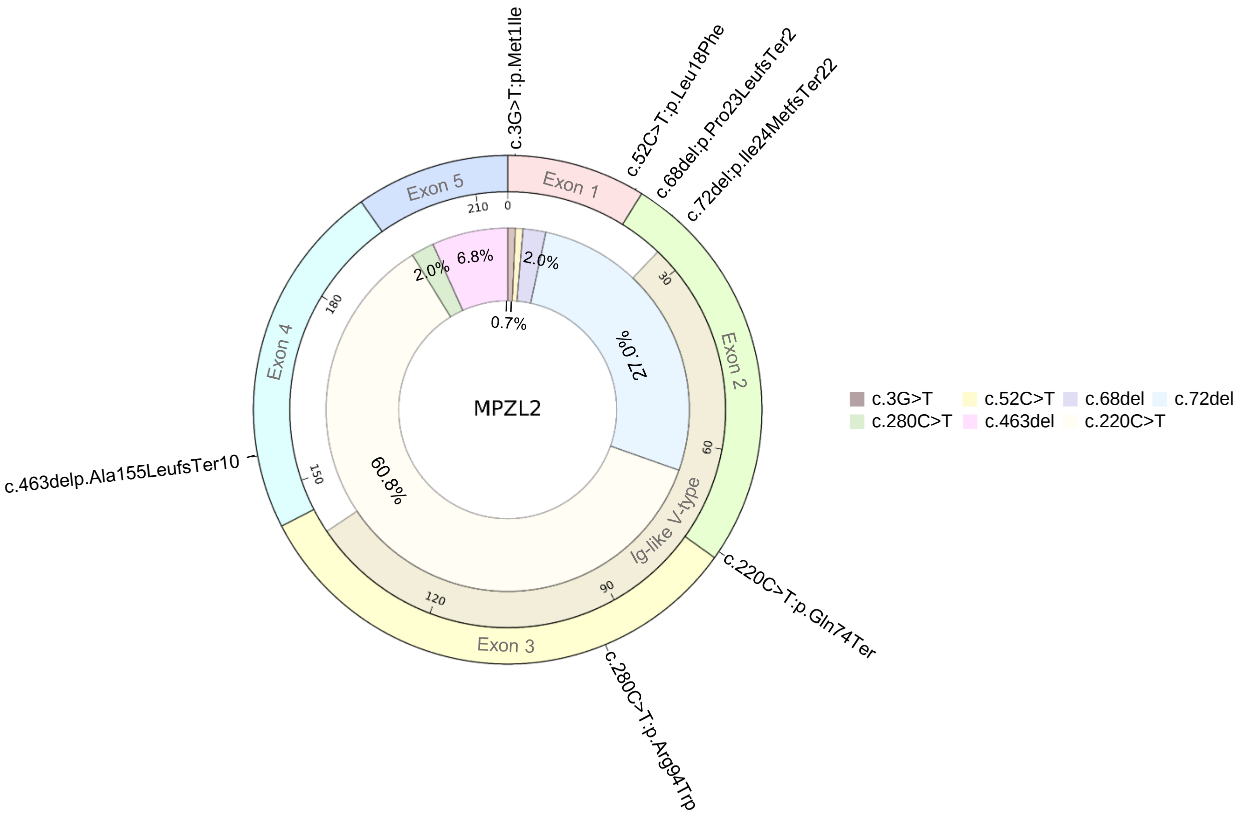


**b**

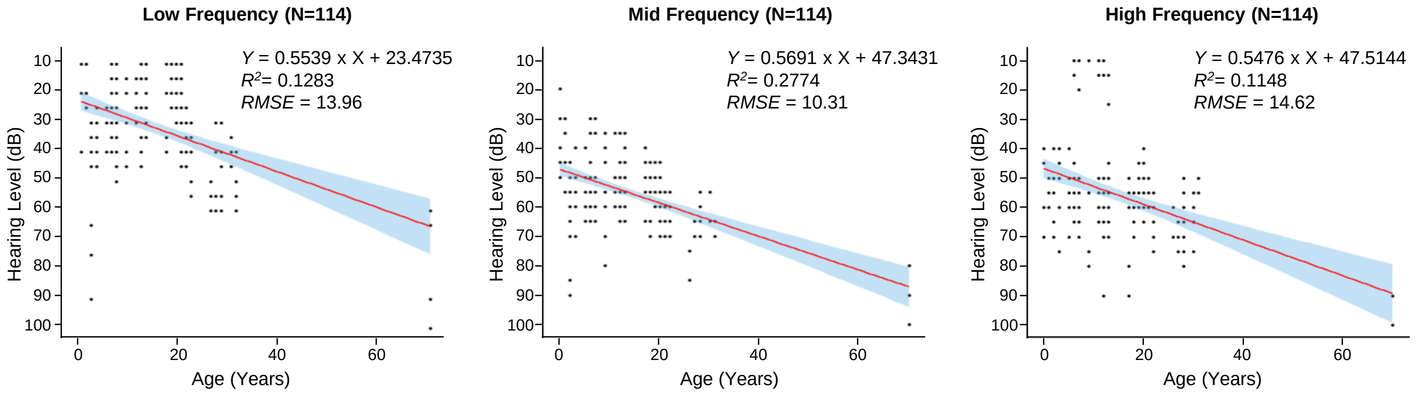

### Extended Data Fig.2

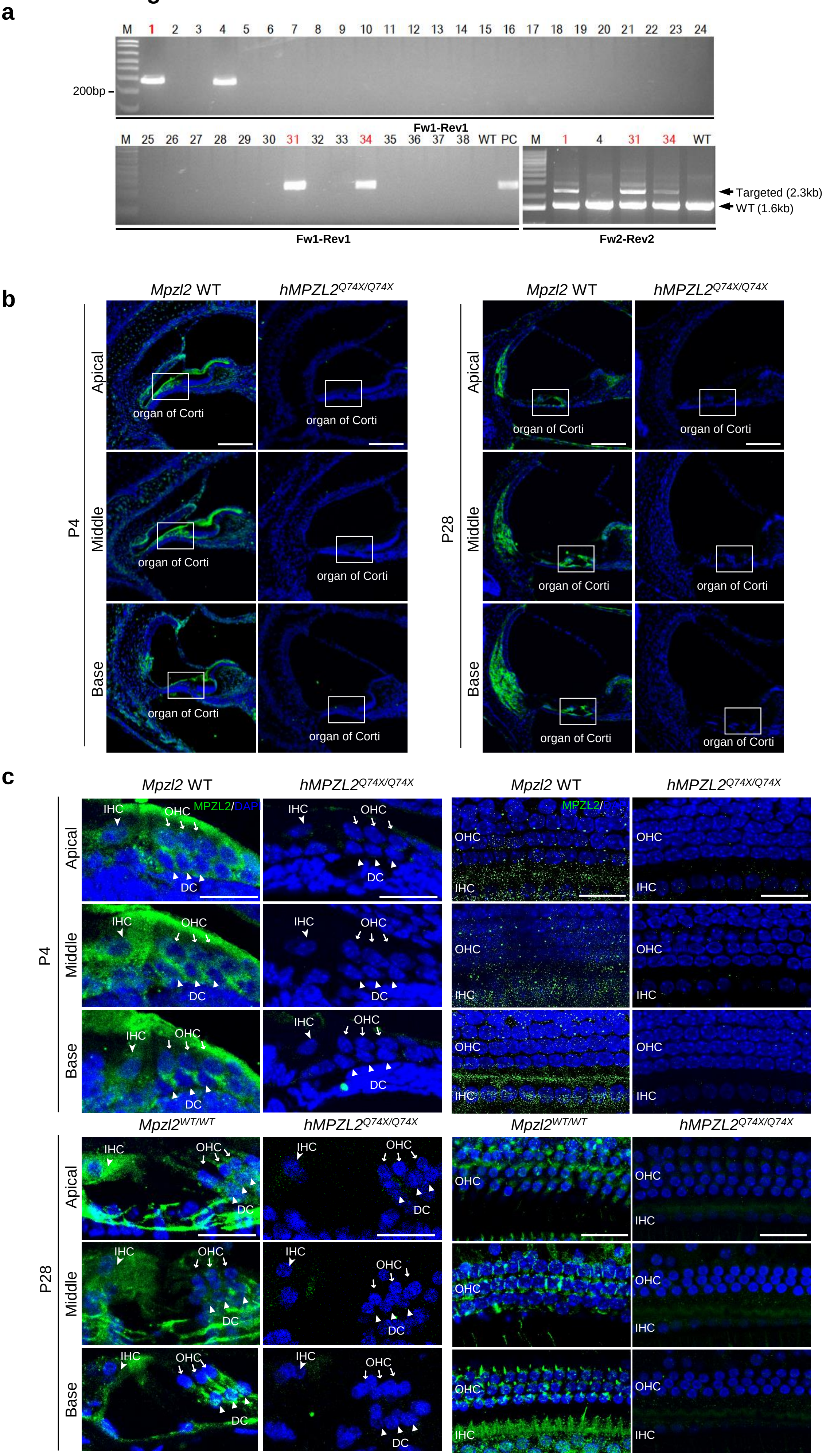

### Extended Data Fig.3

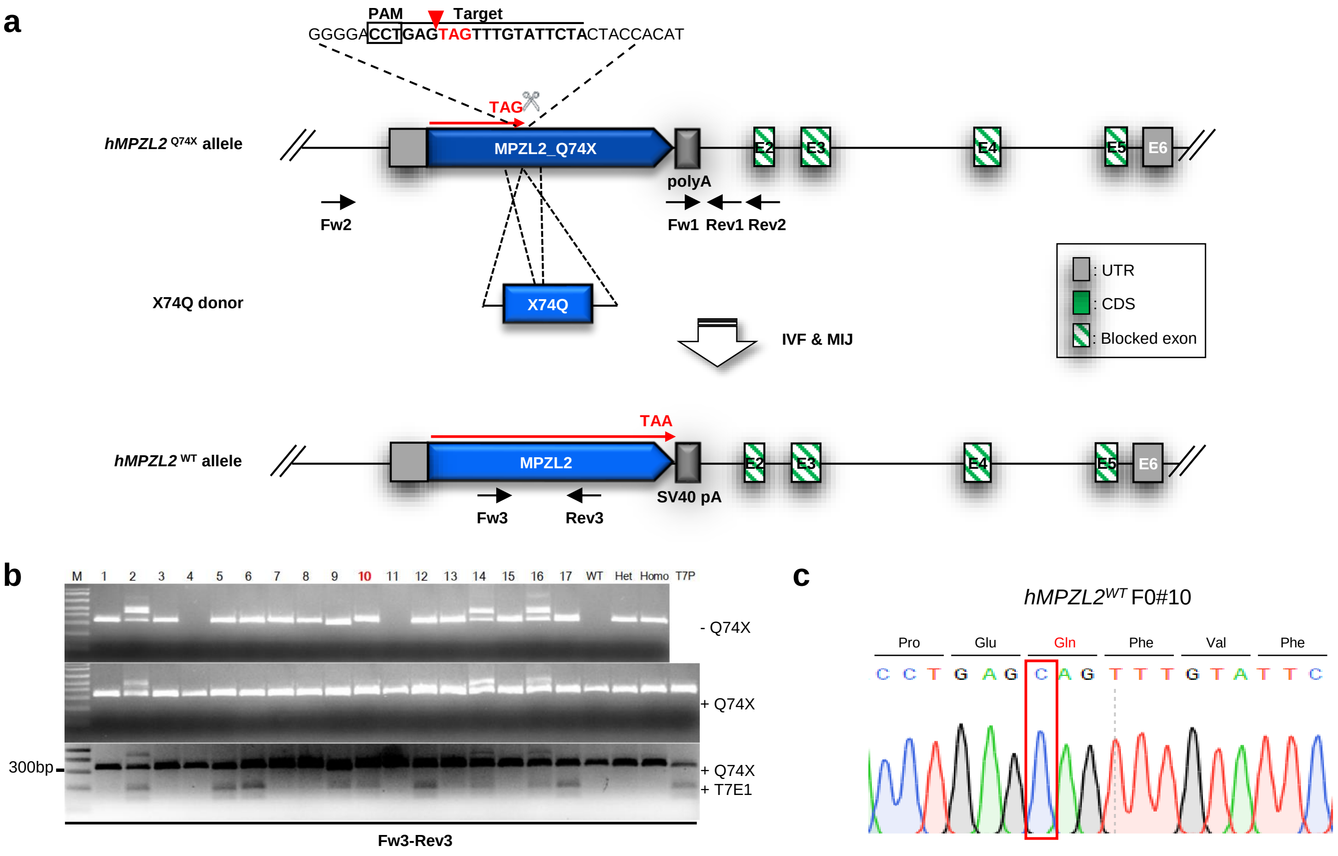

#### Extended Data Fig.4

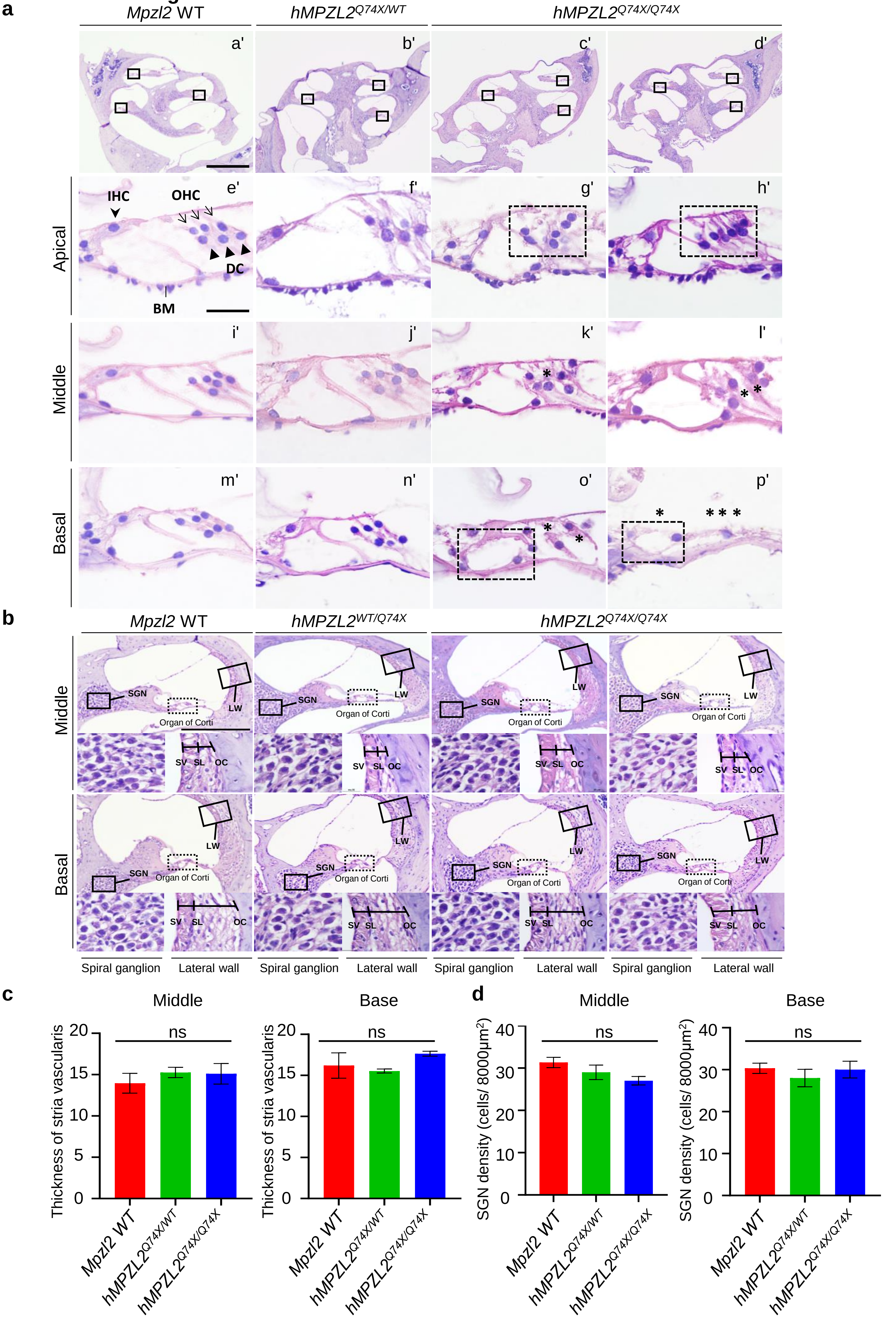

**Extended Data Fig.5**

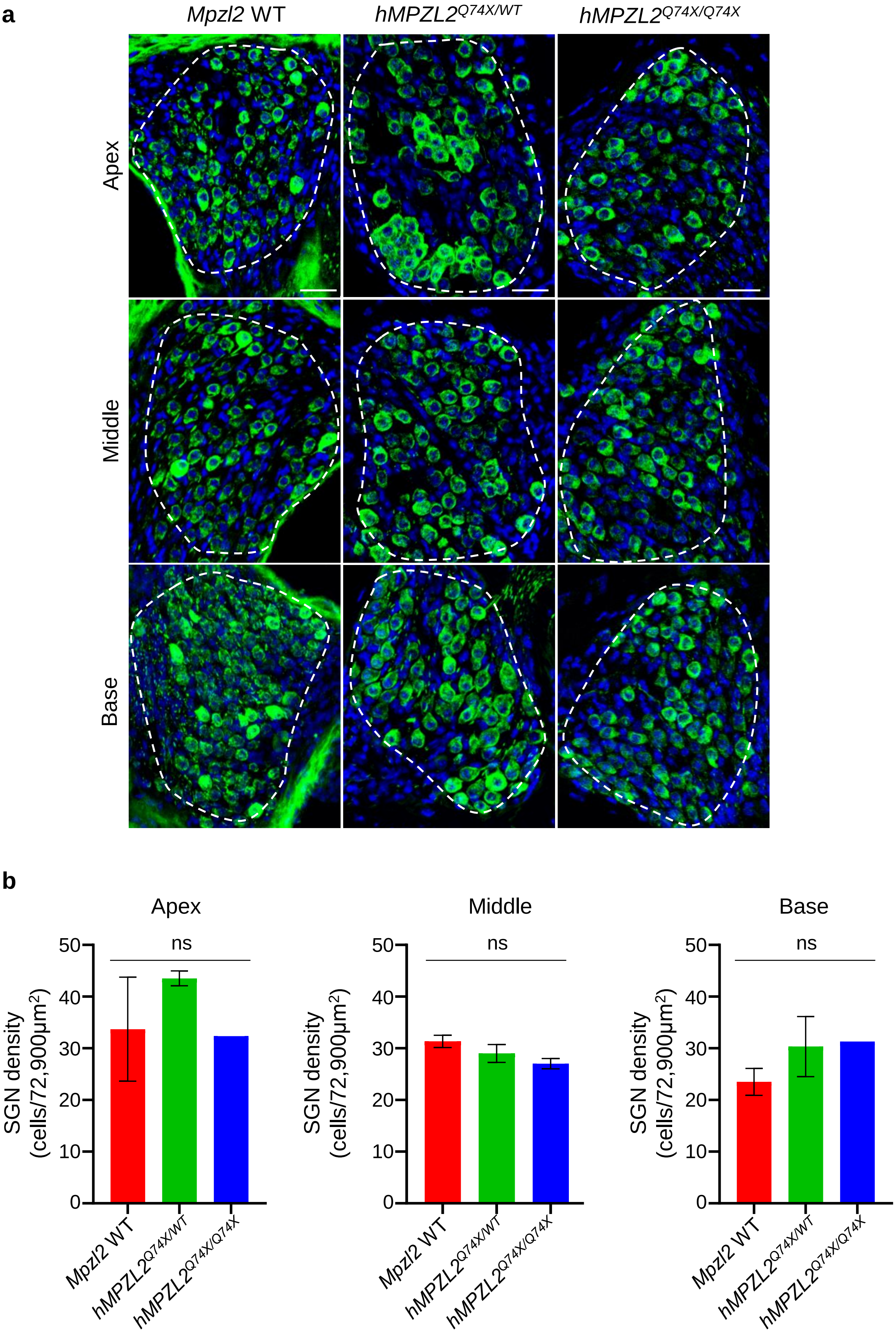

#### Extended Data Fig.6

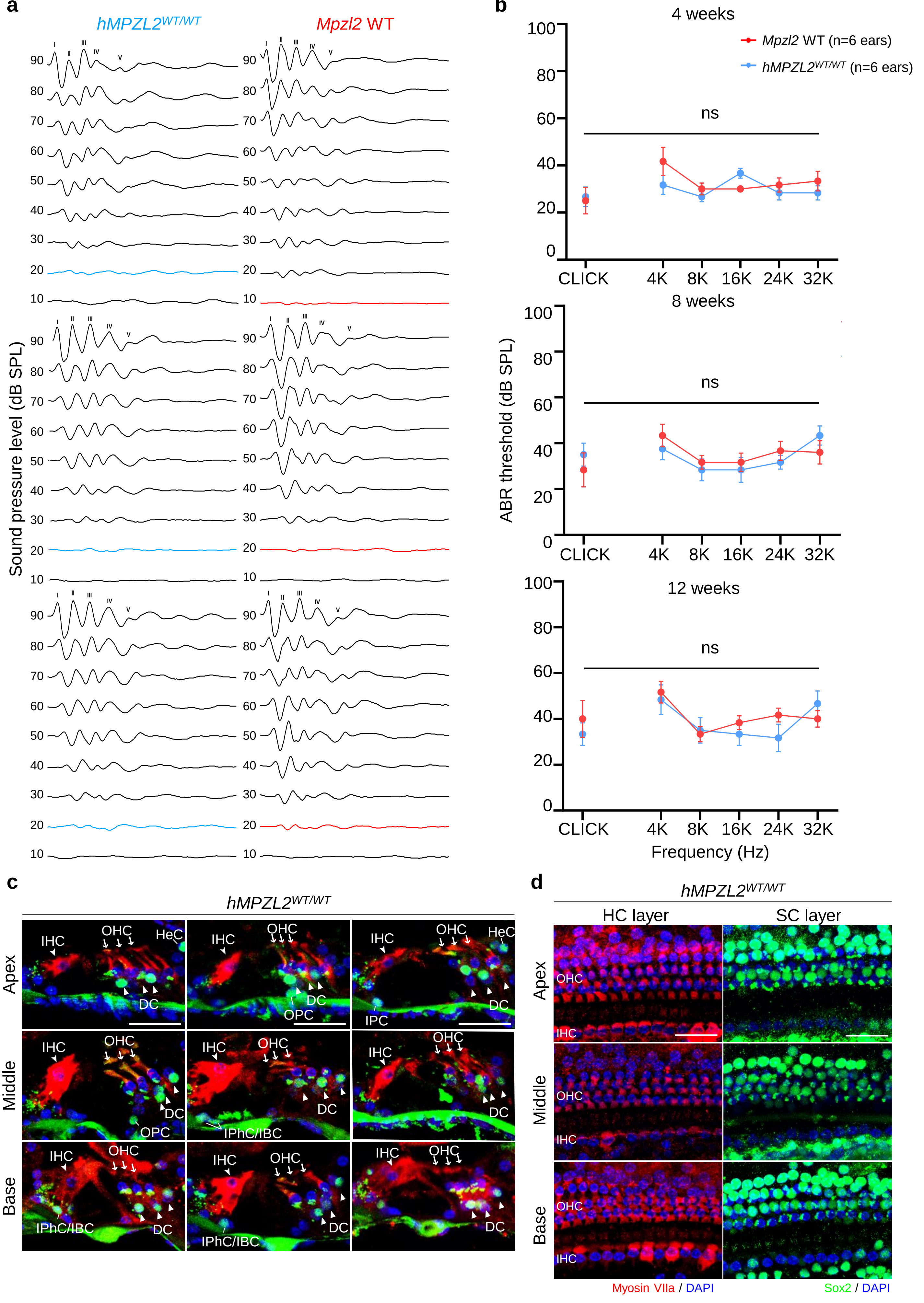

**Extended Data Fig.7**

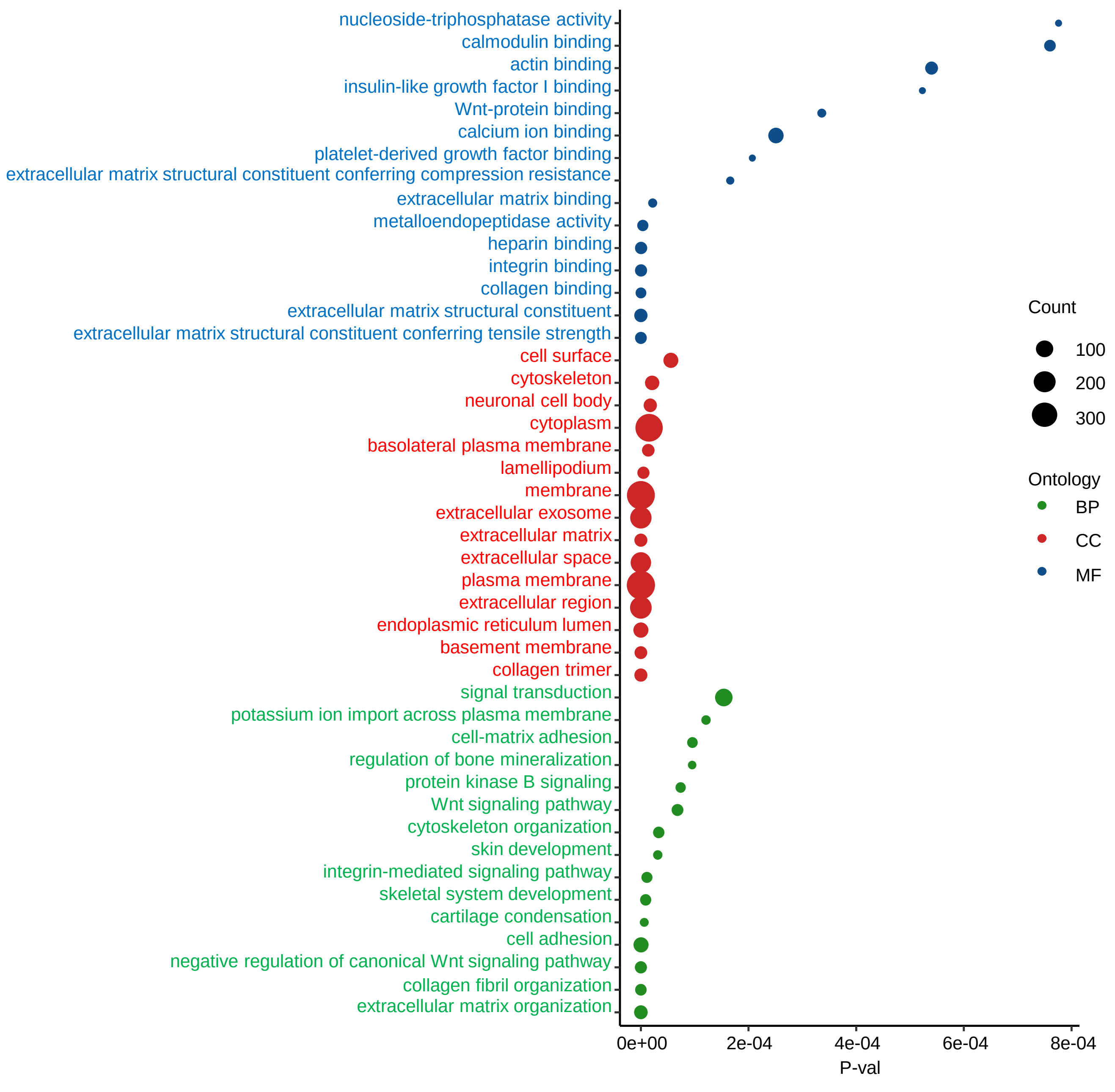

#### Extended Data Fig.8

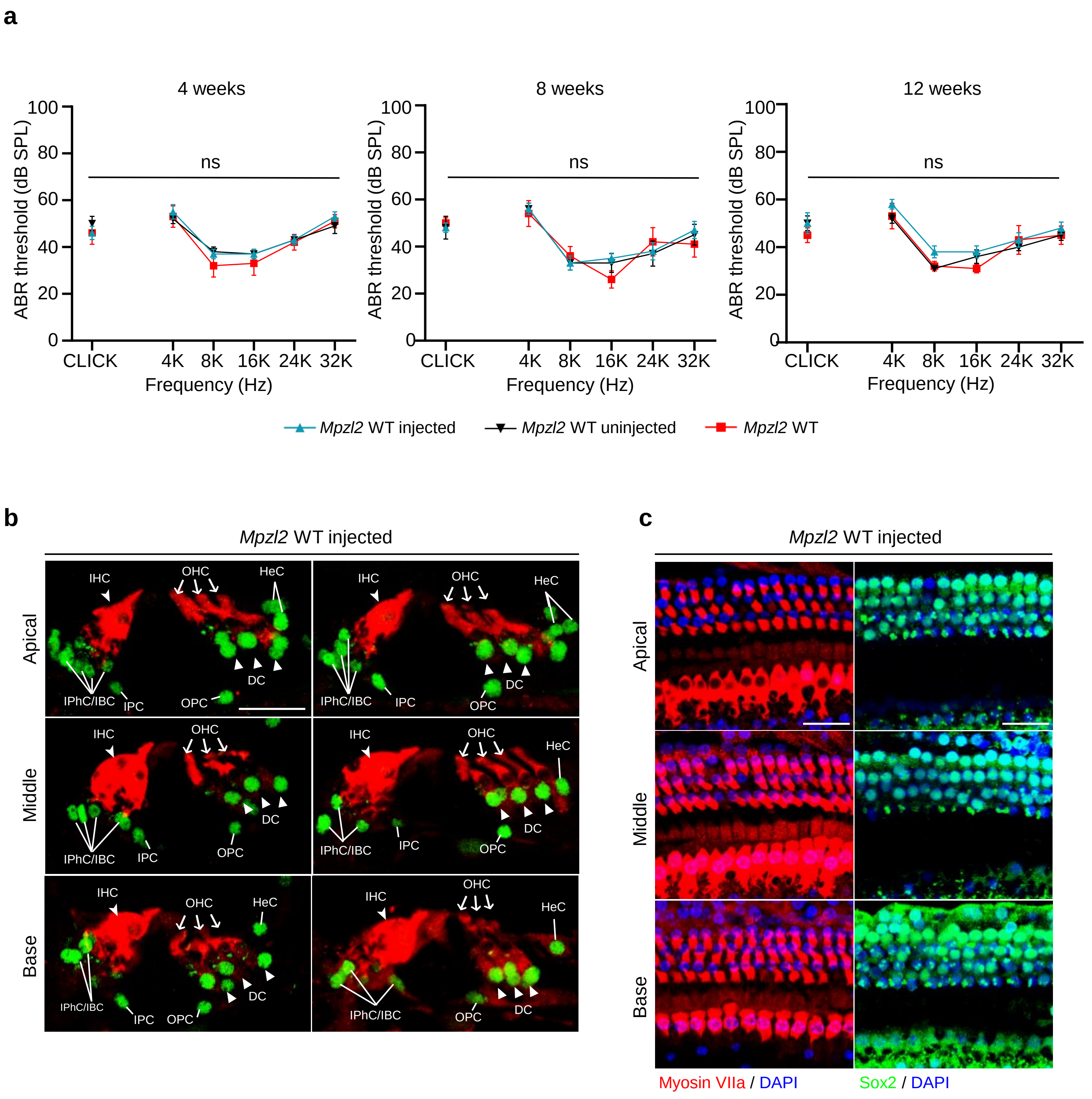

#### Extended Data Fig.9

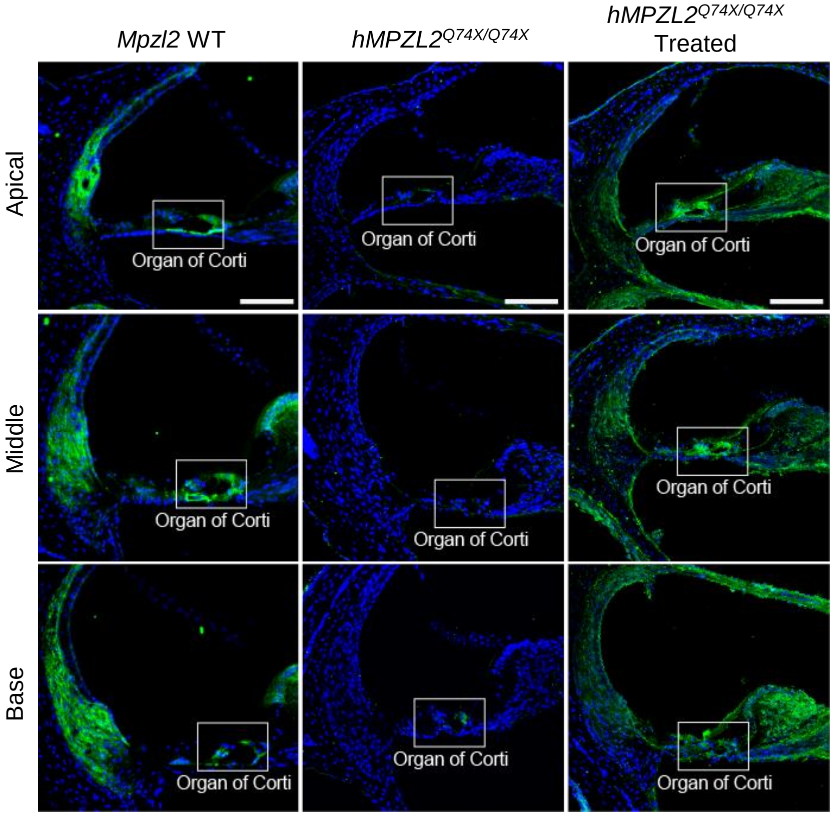

#### Extended Data Fig.10

**a**

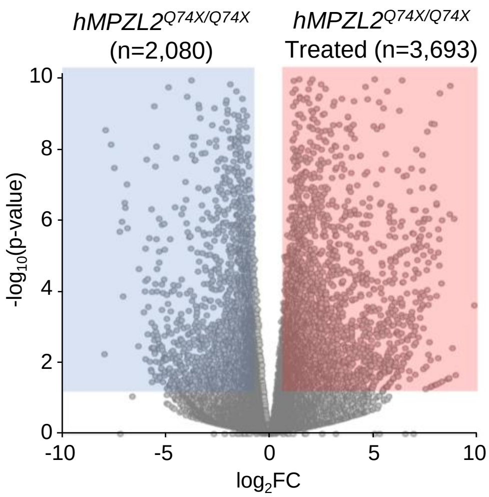


**b**

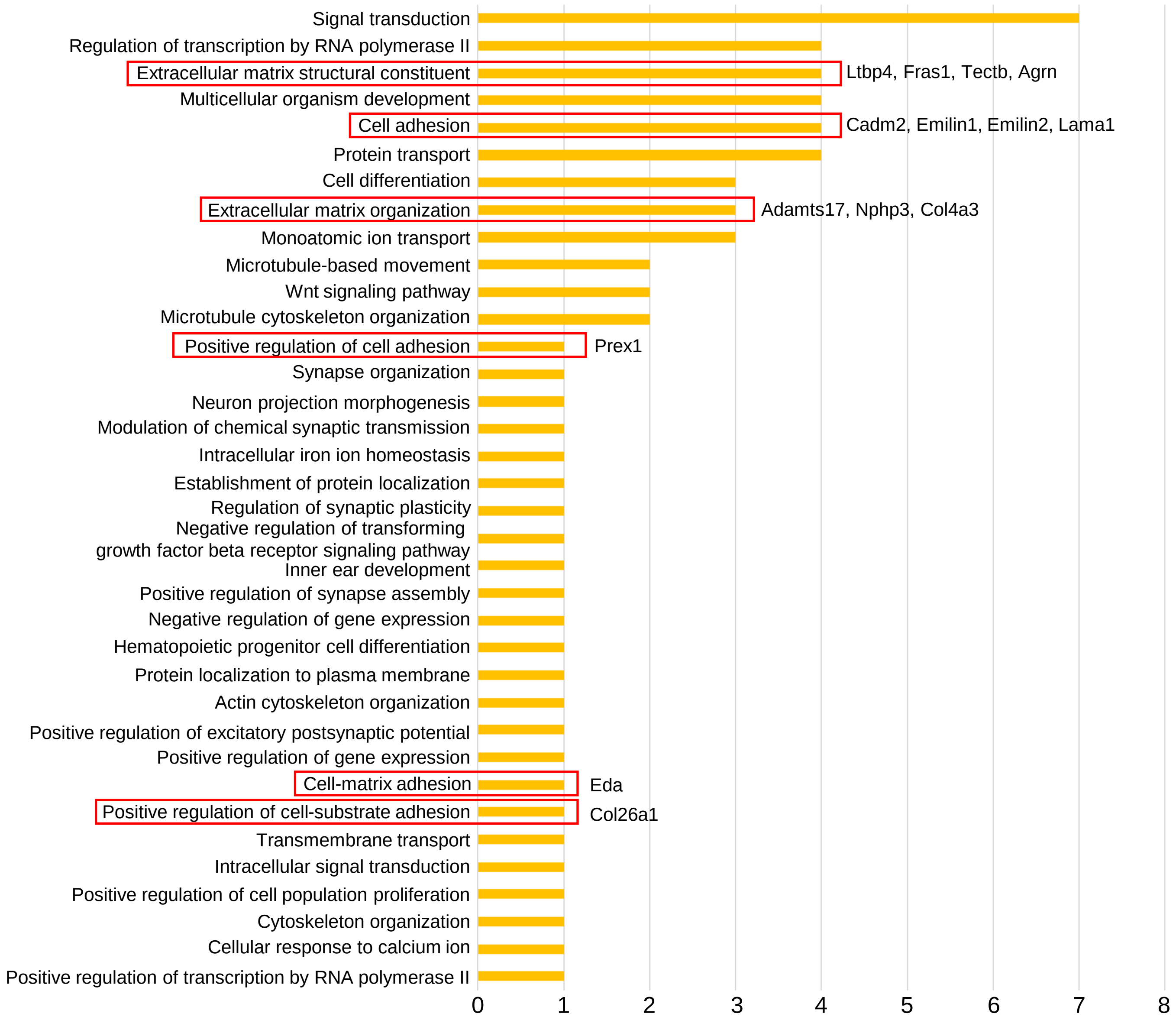
