## Supplementary Table 1 for "PAM-flexible adenine base editing rescues hearing loss in a humanized *MPZL2* mouse model harboring an East Asian founder mutation"

**Supplementary Table 1. Summary of genetic diagnostic steps, identified causative mutations, their pathogenicity, and novelty in 155 genetically diagnosed patients with nonsyndromic, symmetric, mild-to-moderate sensorineural hearing loss.**

| **No.** | **Genetic Diagnostic Step** | | | **Causative Mutations, Pathogenicity, and Novelty** | | | | | | |
| --- | --- | --- | --- | --- | --- | --- | --- | --- | --- | --- |
|  | **WES** | **MLPA/**  **mtDNA** | **WGS** | **Gene** | **NM_Number** | **Mutation 1** | **Mutation 2** | **ACMG-MAP 2018^a^**  **(Mutation 1/Mutation 2)** | **Zygosity** | **Novel^b^**  **(Mutation 1/Mutation 2)** |
| 1 | v |  |  | GJB2 | NM_004004.6 | c.109G>A:p.Val37Ile | c.416G>A:p.Ser139Asn | P / P | comp.het | Reported / Reported |
| 2 | v |  |  | GJB2 | NM_004004.6 | c.235del:p.Leu79CysfsTer3 | c.257C>G:p.Thr86Arg | P / P | comp.het | Reported / Reported |
| 3 | v |  |  | GJB2 | NM_004004.6 | c.109G>A:p.Val37Ile | c.427C>T:p.Arg143Trp | P / P | comp.het | Reported / Reported |
| 4 | v |  |  | GJB2 | NM_004004.6 | c.109G>A:p.Val37Ile | c.427C>T:p.Arg143Trp | P / P | comp.het | Reported / Reported |
| 5 | v |  |  | GJB2 | NM_004004.6 | c.109G>A:p.Val37Ile | c.427C>T:p.Arg143Trp | P / P | comp.het | Reported / Reported |
| 6 | v |  |  | GJB2 | NM_004004.6 | c.109G>A:p.Val37Ile | c.109G>A:p.Val37Ile | P | homo | Reported |
| 7 | v |  |  | GJB2 | NM_004004.6 | c.109G>A:p.Val37Ile | c.176_191del:p.Gly59AlafsTer18 | P / P | comp.het | Reported / Reported |
| 8 | v |  |  | GJB2 | NM_004004.6 | c.235del:p.Leu79CysfsTer3 | c.235del:p.Leu79CysfsTer3 | P | homo | Reported |
| 9 | v |  |  | GJB2 | NM_004004.6 | c.235del:p.Leu79CysfsTer3 | c.109G>A:p.Val37Ile | P / P | comp.het | Reported / Reported |
| 10 | v |  |  | GJB2 | NM_004004.6 | c.109G>A:p.Val37Ile | c.427C>T:p.Arg143Trp | P / P | comp.het | Reported / Reported |
| 11 | v |  |  | GJB2 | NM_004004.6 | c.235del:p.Leu79CysfsTer3 | c.560_605dup:p.Cys202Ter | P / P | comp.het | Reported / Reported |
| 12 | v |  |  | GJB2 | NM_004004.6 | c.109G>A:p.Val37Ile | c.235del:p.Leu79CysfsTer3 | P / P | comp.het | Reported / Reported |
| 13 | v |  |  | GJB2 | NM_004004.6 | c.109G>A:p.Val37Ile | c.176_191del:p.Gly59AlafsTer18 | P / P | comp.het | Reported / Reported |
| 14 | v |  |  | GJB2 | NM_004004.6 | c.109G>A:p.Val37Ile | c.109G>A:p.Val37Ile | P | homo | Reported |
| 15 | v |  |  | GJB2 | NM_004004.6 | c.109G>A:p.Val37Ile | c.109G>A:p.Val37Ile | P | homo | Reported |
| 16 | v |  |  | GJB2 | NM_004004.6 | c.109G>A:p.Val37Ile | c.109G>A:p.Val37Ile | P | homo | Reported |
| 17 | v |  |  | GJB2 | NM_004004.6 | c.109G>A:p.Val37Ile | c.235del:p.Leu79CysfsTer3 | P / P | comp.het | Reported / Reported |
| 18 | v |  |  | GJB2 | NM_004004.6 | c.109G>A:p.Val37Ile | c.109G>A:p.Val37Ile | P | homo | Reported |
| 19 | v |  |  | GJB2 | NM_004004.6 | c.109G>A:p.Val37Ile | c.139G>T:p.Glu47Ter | P / P | comp.het | Reported / Reported |
| 20 | v |  |  | GJB2 | NM_004004.6 | c.109G>A:p.Val37Ile | c.109G>A:p.Val37Ile | P | homo | Reported |
| 21 | v |  |  | GJB2 | NM_004004.6 | c.560_605dup:p.Cys202Ter | c.235del:p.Leu79CysfsTer3 | P / P | comp.het | Reported / Reported |
| 22 | v |  |  | GJB2 | NM_004004.6 | c.109G>A:p.Val37Ile | c.235del:p.Leu79CysfsTer3 | P / P | comp.het | Reported / Reported |
| 23 | v |  |  | GJB2 | NM_004004.6 | c.109G>A:p.Val37Ile | c.299_300del:p.His100ArgfsTer14 | P / P | comp.het | Reported / Reported |
| 24 | v |  |  | GJB2 | NM_004004.6 | c.109G>A:p.Val37Ile | c.109G>A:p.Val37Ile | P | homo | Reported |
| 25 | v |  |  | GJB2 | NM_004004.6 | c.109G>A:p.Val37Ile | c.109G>A:p.Val37Ile | P | homo | Reported |
| 26 | v |  |  | GJB2 | NM_004004.6 | c.109G>A:p.Val37Ile | c.109G>A:p.Val37Ile | P | homo | Reported |
| 27 | v |  |  | GJB2 | NM_004004.6 | c.109G>A:p.Val37Ile | c.109G>A:p.Val37Ile | P | homo | Reported |
| 28 | v |  |  | GJB2 | NM_004004.6 | c.109G>A:p.Val37Ile | c.109G>A:p.Val37Ile | P | homo | Reported |
| 29 | v |  |  | GJB2 | NM_004004.6 | c.109G>A:p.Val37Ile | c.109G>A:p.Val37Ile | P | homo | Reported |
| 30 | v |  |  | GJB2 | NM_004004.6 | c.109G>A:p.Val37Ile | c.109G>A:p.Val37Ile | P | homo | Reported |
| 31 | v |  |  | GJB2 | NM_004004.6 | c.257C>G:p.Thr86Arg | c.560_605dup:p.Cys202Ter | P / P | comp.het | Reported / Reported |
| 32 | v |  |  | GJB2 | NM_004004.6 | c.109G>A:p.Val37Ile | c.109G>A:p.Val37Ile | P | homo | Reported |
| 33 | v |  |  | GJB2 | NM_004004.6 | c.109G>A:p.Val37Ile | c.109G>A:p.Val37Ile | P | homo | Reported |
| 34 | v |  |  | GJB2 | NM_004004.6 | c.235del:p.Leu79CysfsTer3 | c.235del:p.Leu79CysfsTer3 | P | homo | Reported |
| 35 | v |  |  | GJB2 | NM_004004.6 | c.109G>A:p.Val37Ile | c.299_300del:p.His100ArgfsTer14 | P / P | comp.het | Reported / Reported |
| 36 | v |  |  | GJB2 | NM_004004.6 | c.109G>A:p.Val37Ile | c.109G>A:p.Val37Ile | P | homo | Reported |
| 37 | v |  |  | GJB2 | NM_004004.6 | c.109G>A:p.Val37Ile | c.109G>A:p.Val37Ile | P | homo | Reported |
| 38 | v |  |  | GJB2 | NM_004004.6 | c.389G>C:p.Gly130Ala | c.-22-137T>A:p.? | LP / VUS | comp.het | Reported / Novel |
| 39 | v |  |  | GJB2 | NM_004004.6 | c.235del:p.Leu79CysfsTer3 | c.235del:p.Leu79CysfsTer3 | P | homo | Reported |
| 40 | v |  |  | GJB2 | NM_004004.6 | c.109G>A:p.Val37Ile | c.235del:p.Leu79CysfsTer3 | P / P | comp.het | Reported / Reported |
| 41 | v |  |  | GJB2 | NM_004004.6 | c.109G>A:p.Val37Ile | c.109G>A:p.Val37Ile | P | homo | Reported |
| 42 | v |  |  | GJB2 | NM_004004.6 | c.109G>A:p.Val37Ile | c.109G>A:p.Val37Ile | P | homo | Reported |
| 43 | v |  |  | GJB2 | NM_004004.6 | c.109G>A:p.Val37Ile | c.235del:p.Leu79CysfsTer3 | P / P | comp.het | Reported / Reported |
| 44 | v |  |  | GJB2 | NM_004004.6 | c.235del:p.Leu79CysfsTer3 | c.235del:p.Leu79CysfsTer3 | P | homo | Reported |
| 45 | v |  |  | GJB2 | NM_004004.6 | c.109G>A:p.Val37Ile | c.109G>A:p.Val37Ile | P | homo | Reported |
| 46 | v |  |  | GJB2 | NM_004004.6 | c.109G>A:p.Val37Ile | c.109G>A:p.Val37Ile | P | homo | Reported |
| 47 | v |  |  | GJB2 | NM_004004.6 | c.235del:p.Leu79CysfsTer3 | c.235del:p.Leu79CysfsTer3 | P | homo | Reported |
| 48 | v |  |  | GJB2 | NM_004004.6 | c.235del:p.Leu79CysfsTer3 | c.299_300del:p.His100ArgfsTer14 | P / P | comp.het | Reported / Reported |
| 49 | v |  |  | GJB2 | NM_004004.6 | c.109G>A:p.Val37Ile | c.109G>A:p.Val37Ile | P | homo | Reported |
| 50 | v |  |  | GJB2 | NM_004004.6 | c.109G>A:p.Val37Ile | c.109G>A:p.Val37Ile | P | homo | Reported |
| 51 | v |  |  | GJB2 | NM_004004.6 | c.109G>A:p.Val37Ile | c.109G>A:p.Val37Ile | P | homo | Reported |
| 52 | v |  |  | GJB2 | NM_004004.6 | c.235del:p.Leu79CysfsTer3 | c.176_191del:p.Gly59AlafsTer18 | P / P | comp.het | Reported / Reported |
| 53 | v |  |  | GJB2 | NM_004004.6 | c.235del:p.Leu79CysfsTer3 | c.176_191del:p.Gly59AlafsTer18 | P / P | comp.het | Reported / Reported |
| 54 | v |  |  | GJB2 | NM_004004.6 | c.299_300del:p.His100ArgfsTer14 | c.257C>G:p.Thr86Arg | P / P | comp.het | Reported / Reported |
| 55 | v |  |  | GJB2 | NM_004004.6 | c.109G>A:p.Val37Ile | c.109G>A:p.Val37Ile | P | homo | Reported |
| 56 | v |  |  | GJB2 | NM_004004.6 | c.235del:p.Leu79CysfsTer3 | c.235del:p.Leu79CysfsTer3 | P | homo | Reported |
| 57 | v |  |  | GJB2 | NM_004004.6 | c.109G>A:p.Val37Ile | c.109G>A:p.Val37Ile | P | homo | Reported |
| 58 | v | v |  | STRC | NM_153700.2 | c.583C>T:p.Gln195Ter | Deletion | P / P | comp.het | Reported / Reported |
| 59 | v | v |  | STRC | NM_153700.2 | Deletion | Deletion | P | homo | Reported |
| 60 | v | v |  | STRC | NM_153700.2 | c.583C>T:p.Gln195Ter | Deletion | P / P | comp.het | Reported / Reported |
| 61 | v | v |  | STRC | NM_153700.2 | c.583C>T:p.Gln195Ter | Deletion | P / P | comp.het | Reported / Reported |
| 62 | v | v |  | STRC | NM_153700.2 | Deletion | Deletion | P | homo | Reported |
| 63 | v |  |  | STRC | NM_153700.2 | c.4778C>T:p.Ala1593Val | c.4057C>T:p.Gln1353Ter | LP / P | comp.het | Reported / Reported |
| 64 | v | v |  | STRC | NM_153700.2 | c.4816dup:p.Leu1606ProfsTer25 | Deletion | P / P | comp.het | Novel / Reported |
| 65 | v | v |  | STRC | NM_153700.2 | c.4882T>A:p.Cys1628Ser | Deletion | LP / P | comp.het | Nover / Reported |
| 66 | v | v |  | STRC | NM_153700.2 | c.4057C>T:p.Gln1353Ter | Deletion | P / P | comp.het | Reported / Reported |
| 67 | v | v |  | STRC | NM_153700.2 | Deletion | Deletion | P | homo | Reported |
| 68 | v | v |  | STRC | NM_153700.2 | Deletion | Deletion | P | homo | Reported |
| 69 | v | v |  | STRC | NM_153700.2 | Deletion | Deletion | P | homo | Reported |
| 70 | v | v |  | STRC | NM_153700.2 | Deletion | Deletion | P | homo | Reported |
| 71 | v | v |  | STRC | NM_153700.2 | c.4816dup:p.Leu1606ProfsTer25 | Deletion | P / P | comp.het | Novel / Reported |
| 72 | v | v |  | STRC | NM_153700.2 | Deletion | Deletion | P | homo | Reported |
| 73 | v | v |  | STRC | NM_153700.2 | Deletion | Deletion | P | homo | Reported |
| 74 | v |  |  | STRC | NM_153700.2 | c.4816dup:p.Leu1606ProfsTer25 | c.3357_3364dup:p.Pro1122ArgfsTer22 | P / | comp.het | Novel |
| 75 | v | v |  | STRC | NM_153700.2 | c.4816dup:p.Leu1606ProfsTer25 | Deletion | P / P | comp.het | Novel / Reported |
| 76 | v | v |  | STRC | NM_153700.2 | Deletion | Deletion | P | homo | Reported |
| 77 | v | v |  | STRC | NM_153700.2 | Deletion | Deletion | P | homo | Reported |
| 78 | v | v |  | STRC | NM_153700.2 | Deletion | Deletion | P | homo | Reported |
| 79 | v | v |  | STRC | NM_153700.2 | Deletion | Deletion | P | homo | Reported |
| 80 | v | v |  | STRC | NM_153700.2 | Deletion | Deletion | P | homo | Reported |
| 81 | v | v |  | STRC | NM_153700.2 | Deletion | Deletion | P | homo | Reported |
| 82 | v |  |  | STRC | NM_153700.2 | c.4993G>A:p.Ala1665Thr | c.4993G>A:p.Ala1665Thr | LP | homo | Novel |
| 83 | v | v |  | STRC | NM_153700.2 | Deletion | Deletion | P | homo | Reported |
| 84 | v | v |  | STRC | NM_153700.2 | Deletion | Deletion | P | homo | Reported |
| 85 | v | v |  | STRC | NM_153700.2 | Deletion | Deletion | P | homo | Reported |
| 86 | v |  |  | MPZL2 | NM_005797.4 | c.220C>T:p.Gln74Ter | c.220C>T:p.Gln74Ter | P | homo | Reported |
| 87 | v |  |  | MPZL2 | NM_005797.4 | c.220C>T:p.Gln74Ter | c.220C>T:p.Gln74Ter | P | homo | Reported |
| 88 | v |  |  | MPZL2 | NM_005797.4 | c.220C>T:p.Gln74Ter | c.220C>T:p.Gln74Ter | P | homo | Reported |
| 89 | v |  |  | MPZL2 | NM_005797.4 | c.220C>T:p.Gln74Ter | c.436del:p.Ala155LeufsTer10 | P / P | comp.het | Reported / Reported |
| 90 | v |  |  | MPZL2 | NM_005797.4 | c.220C>T:p.Gln74Ter | c.220C>T:p.Gln74Ter | P | homo | Reported |
| 91 | v |  |  | MPZL2 | NM_005797.4 | c.220C>T:p.Gln74Ter | c.220C>T:p.Gln74Ter | P | homo | Reported |
| 92 | v |  |  | MPZL2 | NM_005797.4 | c.220C>T:p.Gln74Ter | c.220C>T:p.Gln74Ter | P | homo | Reported |
| 93 | v |  |  | MPZL2 | NM_005797.4 | c.220C>T:p.Gln74Ter | c.220C>T:p.Gln74Ter | P | homo | Reported |
| 94 | v |  |  | MPZL2 | NM_005797.4 | c.220C>T:p.Gln74Ter | c.220C>T:p.Gln74Ter | P | homo | Reported |
| 95 | v |  |  | MPZL2 | NM_005797.4 | c.220C>T:p.Gln74Ter | c.220C>T:p.Gln74Ter | P | homo | Reported |
| 96 | v |  |  | MPZL2 | NM_005797.4 | c.220C>T:p.Gln74Ter | c.220C>T:p.Gln74Ter | P | homo | Reported |
| 97 | v |  |  | MPZL2 | NM_005797.4 | c.220C>T:p.Gln74Ter | c.220C>T:p.Gln74Ter | P | homo | Reported |
| 98 | v |  |  | MPZL2 | NM_005797.4 | c.220C>T:p.Gln74Ter | c.220C>T:p.Gln74Ter | P | homo | Reported |
| 99 | v |  |  | MPZL2 | NM_005797.4 | c.220C>T:p.Gln74Ter | c.463del:p.Ala155LeufsTer10 | P / P | comp.het | Reported / Reported |
| 100 | v |  |  | USH2A | NM_206933.4 | c.8559-2A>G:p.? | c.11156G>A:p.Arg3719His | P / P | comp.het | Reported /Reported |
| 101 | v | v | v | USH2A | NM_206933.4 | c.10712C>T:p.Thr3571Met | c.7120+1475A>G:p.? | P / LP | comp.het | Reported /Novel |
| 102 | v |  |  | USH2A | NM_206933.4 | c.2802T>G:p.Cys934Trp | c.4858C>T:p.Gln1620Ter | LP / P | comp.het | Reported /Reported |
| 103 | v | v | v | USH2A | NM_206933.4 | c.14835del:p.Val4946TrpfsTer4 | c.14134-3169A>G:p.? | P / LP | comp.het | Reported / Reported |
| 104 | v |  |  | USH2A | NM_206933.4 | c.11156G>A:p.Arg3719His | c.8559-2A>G:p.? | P / P | comp.het | Reported / Reported |
| 105 | v |  |  | USH2A | NM_206933.4 | c.8559-2A>G:p.? | c.11389+3A>T:p.? | P / P | comp.het | Reported / Reported |
| 106 | v |  |  | USH2A | NM_206933.4 | c.8559-2A>G:p.? | c.8559-2A>G:p.? | P | homo | Reported |
| 107 | v |  |  | USH2A | NM_206933.4 | c.14287G>A:p.Gly4763Arg | c.14791+5G>A:p.? | LP / VUS | comp.het | Reported / Novel |
| 108 | v |  |  | USH2A | NM_206933.4 | c.6485+5G>A:p.? | c.8559-2A>G:p.? | P / P | comp.het | Reported / Reported |
| 109 | v |  |  | OTOGL | NM_001378609.3 | c.2860C>T:p.Arg954Ter | c.6162_6163del:p.Ala2055ArgfsTer21 | P / P | comp.het | Reported / Novel |
| 110 | v |  |  | OTOGL | NM_001378609.3 | c.6162_6163del:p.Ala2055ArgfsTer21 | c.6162_6163del:p.Ala2055ArgfsTer21 | P | homo | Novel |
| 111 | v |  |  | OTOGL | NM_001378609.3 | c.3058A>T:p.Lys1020Ter | c.6162_6163del:p.Ala2055ArgfsTer21 | LP / P | comp.het | Novel / Novel |
| 112 | v |  |  | OTOGL | NM_001378609.3 | c.5743C>T:p.Arg1915Ter | c.6162_6163del:p.Ala2055ArgfsTer21 | LP / P | comp.het | Reported / Novel |
| 113 | v |  |  | OTOGL | NM_001378609.3 | c.6409G>T:p.Asp2137Tyr | c.2833C>T:p.Pro945Ser | P / P | comp.het | Novel / Novel |
| 114 | v |  |  | OTOGL | NM_001378609.3 | c.6494C>A:p.Ser2165Ter | c.6494C>A:p.Ser2165Ter | LP | homo | Reported |
| 115 | v |  |  | OTOA | NM_144672.4 | c.1176T>G:p.Asp392Glu | c.1765del:p.Gln589ArgfsTer55 | LP/ P | comp.het | Novel / Reported |
| 116 | v |  |  | OTOA | NM_144672.4 | c.1765del:p.Gln589ArgfsTer55 | c.3366G>A:p.Trp1122Ter | P / P | comp.het | Reported / Novel |
| 117 | v | v |  | OTOA | NM_144672.4 | Deletion | Deletion | P | homo | Reported |
| 118 | v | v |  | OTOA | NM_144672.4 | Deletion | Deletion | P | homo | Reported |
| 119 | v |  |  | GSDME | NM_001127453.1 | c.1183+4A>G:p.? |  | P | het | Reported |
| 120 | v |  |  | GSDME | NM_001127453.1 | c.991-15_991-13del |  | P | het | Reported |
| 121 | v |  |  | GSDME | NM_001127453.1 | c.991-15_991-13del |  | P | het | Reported |
| 122 | v |  |  | MYH14 | NM_001145809.2 | c.2375A>T:p.Leu792Gln |  | VUS | het | Novel |
| 123 | v |  |  | MYH14 | NM_001145809.2 | c.6008del:p.Arg2003HisfsTer2 |  | P | het | Novel |
| 124 | v |  |  | MYH14 | NM_001145809.2 | c.5990del:p.Thr1997ArgfsTer8 |  | P | het | Reported |
| 125 | v |  |  | MYO7A | NM_000260.4 | c.2558G>A:p.Arg853His |  | LP | het | Reported |
| 126 | v |  |  | MYO7A | NM_000260.4 | c.2558G>A:p.Arg853His |  | LP | het | Reported |
| 127 | v |  |  | MYO7A | NM_000260.4 | c.2011G>A:p.Gly671Ser |  | LP | het | Reported |
| 128 | v |  |  | OTOG | NM_001277269.2 | c.330C>G:p.Tyr110Ter | c.330C>G:p.Tyr110Ter | P | homo | Reported |
| 129 | v |  |  | OTOG | NM_001277269.2 | c.330C>G:p.Tyr110Ter | c.330C>G:p.Tyr110Ter | P | homo | Reported |
| 130 | v |  |  | OTOG | NM_001277269.2 | c.330C>G:p.Tyr110Ter | c.704del:p.Leu235ArgfsTer39 | P / P | comp.het | Reported / Novel |
| 131 | v |  |  | SIX1 | NM_005982.4 | c.397_399del:p.Glu133del |  | P | het | Reported |
| 132 | v |  |  | SIX1 | NM_005982.4 | c.386_391del:p.Tyr129_Cys130del |  | P | het | Reported |
| 133 | v |  |  | SIX1 | NM_005982.4 | c.376_378del:p.Glu126del |  | LP | het | Novel |
| 134 | v |  |  | EYA4 | NM_001301013.1 | c.1777C>T:p.Arg593Ter |  | P | het | Reported |
| 135 | v |  | v | EYA4 | NM_001301013.1 | Deletion |  | P | het | Novel |
| 136 | v |  |  | KCNQ4 | NM_004700.4 | c.961G>A:Gly321Ser |  | P | het | Reported |
| 137 | v |  |  | KCNQ4 | NM_004700.4 | c.853G>A:Gly285Ser |  | P | het | Reported |
| 138 | v |  |  | MYO6 | NM_004999.4 | c.2354_2390del:p.Asn785SerfsTer10 |  | LP | het | Novel |
| 139 | v |  |  | MYO6 | NM_004999.4 | c.2751dup:p.Gln918ThrfsTer24 |  | P | het | Reported |
| 140 | v |  |  | TECTA | NM_005422.4 | c.494C>T:p.Thr165Ile |  | LP | het | Novel |
| 141 | v |  |  | TECTA | NM_005422.4 | c.5597C>T:p.Thr1866Met |  | P | het | Reported |
| 142 | v |  |  | WFS1 | NM_006005.3 | c.2389G>A:p.Asp797Asn |  | LP | het | Reported |
| 143 | v |  |  | WFS1 | NM_006005.3 | c.2530G>A:p.Ala844Thr |  | LP | het | Reported |
| 144 | v | v | v | CLCNKA_CLCNKB | NM_004070.4 | c.778C>T:p.Gln260Ter | g.[16032250_16046349del] | P / P | comp.het | Reported / Novel |
| 145 | v |  |  | GATA3 | NM_001002295.2 | c.1047C>G:p.His349Gln |  | LP | het | Novel |
| 146 | v |  |  | ADGRV1 | NM_032119.4 | c.7406G>A;p.Trp2469Ter | c.7406G>A;p.Trp2469Ter | P | comp.het | Reported |
| 147 | v |  |  | GRHL2 | NM_024915.4 | c.1763+2T>C:p.? |  | LP | het | Novel |
| 148 | v | v |  | MT-RNR1 | - | m.1555A>G |  | P | hetero-  plasmy | Reported |
| 149 | v |  |  | MYO15A | NM_032119.4 | c.10250_10252del:p.Ser3417del | c.10263C>G:p.Ile3421Met | LP / LP | comp.get | Reported / Reported |
| 150 | v |  |  | PRPS1 | NM_002764.4 | c.259G>A:p.Ala87Thr |  | LP | hemi | Reported |
| 151 | v |  |  | SCD5 | NM_001037582.3 | c.173G>A:p.Trp158Ter |  | VUS | het | Novel |
| 152 | v |  |  | SLC17A8 | NM_139319.3 | c.449dup:p.Ser151PhefsTer7 |  | P | het | Novel |
| 153 | v |  |  | SLC26A5 | NM_1989993.3 | c.1222G>A:p.Gly408Arg |  | LP | het | Novel |
| 154 | v |  |  | SMPX | NM_014332.3 | c.46-2A>G:p.? |  | LP | hemi | Novel |
| 155 | v |  |  | TMC1 | NM_138691.3 | c.1444T>C:p.Trp482Arg |  | LP | het | Reported |

Abbreviations: WES, whole-exome sequencing; MLPA, Multiplex Ligation-dependent Probe Amplification; WGS, whole-genome sequencing; P, pathogenic; LP, likely pathogenic; VUS, uncertain significance of variant; het, heterozygote; homo, homozygote; hemi, hemizygote; comp.het, compound heterozygote; homo, homozygote; ACMG-AMP, American College of Medical Genetics and Genomics and the Association for Molecular Pathology.

Note^a^: The pathogenicity of identified mutations was classified according to ACMG-AMP2018 guideline.

Note^b^: The presence of novel mutations was determined using the Clinvar database (<https://www.ncbi.nlm.nih.gov/clinvar/>) and the Litvar2 database (<https://www.ncbi.nlm.nih.gov/research/litvar2/>) .
